# Mapping vestibular and visual contributions to angular head velocity tuning in the cortex

**DOI:** 10.1101/2021.04.29.441624

**Authors:** Eivind Hennestad, Aree Witoelar, Anna Chambers, Koen Vervaeke

## Abstract

Neurons that signal the direction and angular velocity of head movements (AHV cells) are critically important to process visual and spatial information. However, it has been challenging to isolate the sensory modality that drives them and to comprehensively map their cortical distribution. To address this, we developed a method that enables rotating awake, head-fixed mice under a two-photon microscope in a visual environment. Starting in layer 2/3 of the retrosplenial cortex, a key area for vision and navigation, we found that a significant fraction of rotation sensitive neurons report AHV. These tuning properties depend on vestibular input because they persist in darkness and are reduced when replaying visual flow to stationary animals. When mapping the spatial extent, we found AHV cells in all cortical areas that we explored, including motor, somatosensory, visual and posterior parietal cortex. Notably, the vestibular and visual contributions to AHV are area dependent. Thus, many cortical circuits have access to AHV, enabling a diverse integration with sensorimotor and cognitive information.

## Introduction

Neural circuits have access to head motion information through a population of neurons whose firing rate is correlated with angular head velocity (AHV) (Bassett and Taube, 2001; Büttner and Buettner, 1978; Sharp et al., 2001; Turner Evans et al., 2017). Beside their well-known importance for eye movements, these cells also enable operations essential for perception and cognition (Angelaki and Cullen, 2008; Cullen, 2019). For example, all moving organisms require self-motion information to determine whether a changing visual stimulus is due to their own movements or due to a moving object (Angelaki and Cullen, 2008; Sasaki et al., 2017; Vélez-Fort et al., 2018). In addition, by integrating angular speed signals across short time windows, the brain can estimate displacement to update the animal’s internal representation of orientation mediated by so-called "head direction cells" (Skaggs et al., 1995; Taube et al., 1990). Such diverse operations likely require the integration of AHV, sensory and motor information at multiple levels along the hierarchy of brain organization. However, while AHV tuning is well characterized in the brain stem and cerebellum, comparatively little is known about AHV tuning in the cortex.

Unlike tuning properties in the visual and auditory system, AHV tuning can depend on multiple sensory modalities. While thought to rely primarily on head motion signals from the vestibular organs, AHV tuning can also rely on neck motor commands (efference copies) (Guitchounts et al., 2020; Medrea and Cullen, 2013; Miall and Wolpert, 1996; Roy and Cullen, 2001), proprioceptive feedback signals from the neck (Gdowski and McCrea, 2000; Medrea and Cullen, 2013; Mergner et al., 1997) and eye muscles (Sikes et al., 1988), and visual flow (Waespe and Henn, 1977). Adding to this complexity, AHV signals can be entangled with linear speed, body posture, and place signals (Cho and Sharp, 2001; McNaughton et al., 1983; Mimica et al., 2018; Sharp, 1996; Wilber et al., 2014). Nevertheless, influential theories about neuronal coding require us to determine the exact modality. For example, models (Cullen and Taube, 2017; Hulse and Jayaraman, 2019; Skaggs et al., 1995) and data (Stackman et al., 2002; Valerio and Taube, 2016) predict that AHV-tuning driven by the vestibular organs plays a critical role in generating head direction tuning. However, because motor commands and the ensuing vestibular and other sensory feedback signals occur quasi-simultaneously, their individual contribution to AHV tuning is difficult to disentangle. Therefore, to characterize AHV tuning in the cortex, we used head-fixed mice that are passively rotated, enhancing experimental control and facilitating the isolation of different modalities.

Recent work in primary visual cortex of the rodent shows that vestibular input plays a central role in AHV tuning and is thought to be conveyed via direct projections from the retrosplenial cortex (RSC) (Bouvier et al., 2020; Vélez-Fort et al., 2018). The RSC is well-positioned to convey head motion information because it plays a key role in vision-based navigation (Vann et al., 2009). About 22 % of RSC neurons are selective for left or right turns during a navigation task (Alexander and Nitz, 2015), and 5-10 % of neurons report heading direction (Alexander and Nitz, 2015; Chen et al., 1994; Cho and Sharp, 2001; Jacob et al., 2016). There are also reports of RSC neurons tuned to AHV in freely moving rodents (Cho and Sharp, 2001), but whether these are driven by vestibular input or by other sensory or motor input is not known. In rodents, it is also not clear whether AHV tuning is limited to specific cortical areas. Stimulation of the vestibular nerve evokes widespread activation of cortical circuits (Lopez et al., 2011; Rancz et al., 2015), but this method does not reveal whether neurons are tuned to angular direction and speed. Therefore, how AHV is represented in RSC, and whether AHV tuning in the cortex is area-specific, remains to be determined.

The lack of knowledge about vestibular representations in the cortex stems, in part, from the fundamental obstacle that the animal’s head needs to move. This has limited the use of powerful methods to study neural circuits. Two-photon microscopy and genetically encoded Ca^2+^ indicators (Chen et al., 2013) enable monitoring the activity of thousands of neurons and have provided tremendous insight into cortical function. However, two-photon microscopy has been limited to stationary animals. Therefore, to overcome this technical limitation, we developed a setup to perform large-scale neuronal recordings from mice rotated under a two-photon microscope. This enables manipulating vestibular and visual stimuli under well-controlled conditions.

Using this method, we found that a significant fraction of neurons in layer 2/3 of RSC are tuned to the direction and speed of angular head movements. We show these tuning properties persist when rotating the animals in darkness and are generally consistent with the hypothesis that AHV tuning is vestibular-dependent. Strikingly, we discovered that AHV tuning is not limited to RSC. By mapping large parts of the cortex, we found that a similar fraction of neurons is tuned to AHV in all cortical areas explored, including visual, somatosensory, motor and posterior parietal cortex. Finally, the contribution of vestibular and visual input to AHV tuning was distinct between areas. This shows that AHV tuning is more widespread in the cortex than previously thought and that the underlying sensory modality is area dependent.

## Results

### Turn-selective neurons in the retrosplenial cortex

How do neurons in retrosplenial cortex (RSC) respond to angular head motion? To control the velocity of head movements, we positioned awake but head-restrained mice in the centre of a motorized platform that enables rotation along the horizontal plane (Figure 1A). A circular enclosure with visual cues surrounded the platform (Figure 1A-B, see Methods). This arena could be rotated independently to determine whether motion-sensitive neurons depend on vestibular input or visual flow. We also performed pupil tracking to determine whether neurons are tuned to eye movements (Video S1). The mice were rotated clockwise (CW) or counter clockwise (CCW) between three randomly chosen target directions (Figure 1B-C). The speed profile of the motion was trapezoidal, with fast accelerations and decelerations (200 °/s2) such that most of the rotation occurred at a constant speed (45 °/s). We also ensured that the animals’ arousal levels remained constant during the entire session by keeping the mice engaged with rewards in the form of small water drops when facing one of the three target positions (Figure 1B-C).

**Figure 1.**
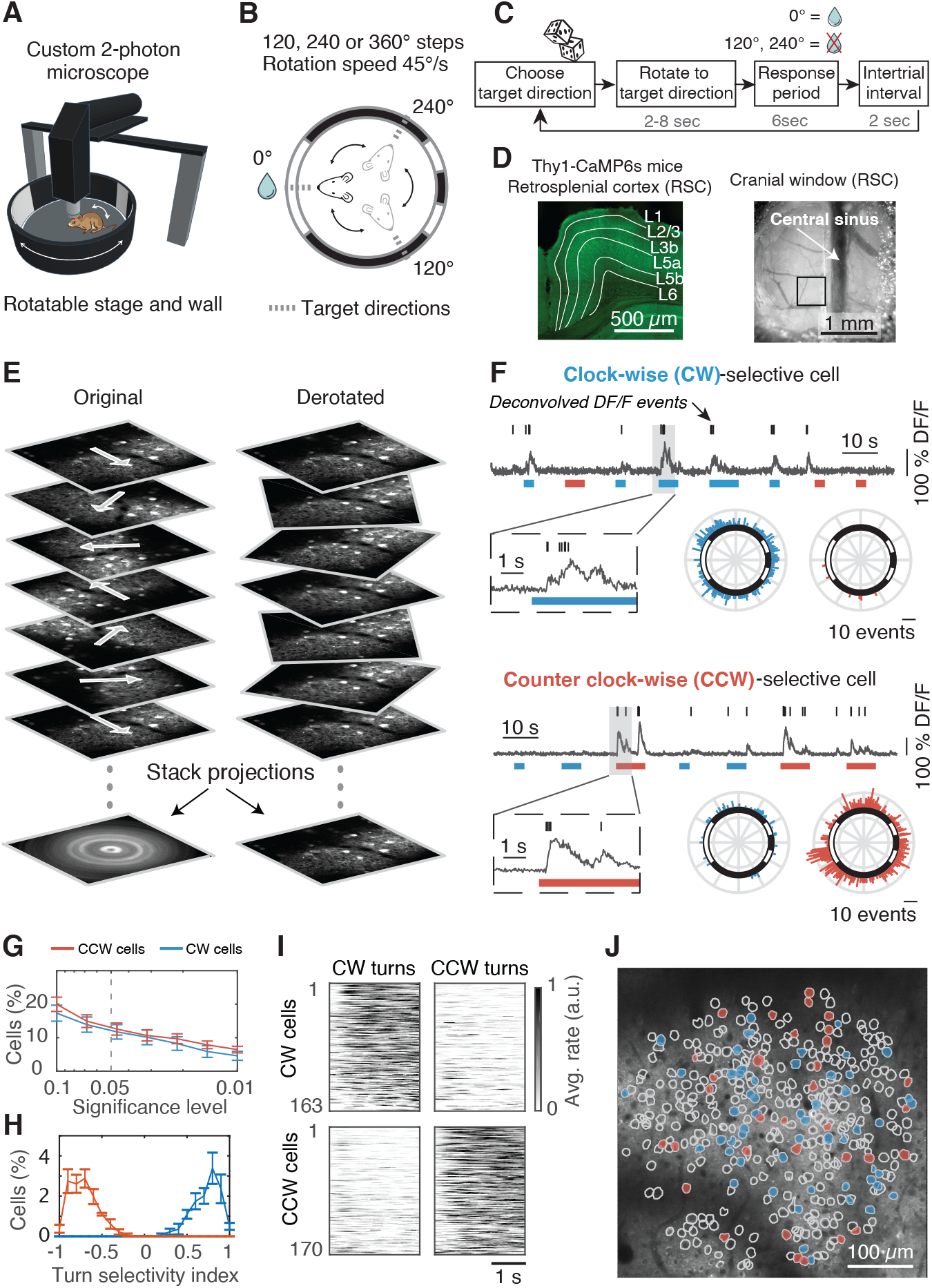
Turn-selective neurons in the retrosplenial cortex. (**A**) Cartoon of head-fixed mouse in an arena with visual cues. The mouse and visual cues can be rotated together or independently while performing two photon microscopy. (**B**) Top-view schematic; the mouse is randomly rotated between three target directions (dashed lines), but only gets a water reward in one direction. The circle represents the arena wall with white cues on a black background. (**C**) Block diagram of the sequence of events for a single trial. (**D**) Left: Confocal fluorescence image of RSC coronal brain slice with cortical layers indicated (Thy1-GCaMP6s/ GP4.3 mice). Right: Top view of cranial window implanted on dorsal (agranular) RSC and centred over the central sinus. Black box shows the typical size and location of an imaging Field of View (FOV). (**E**) Two-photon image sequence of layer 2/3 neurons during rotation experiment (left) and when de-rotated during post-processing (right). (**F**) Example fluorescence traces (fractional change of fluorescence, DF/F) of a clockwise (CW) and counterclockwise (CCW) selective cell. Ticks indicate deconvolved DF/F events. Blue and red bars are CW and CCW rotation trials respectively. Polar histograms show the DF/F events as a function of direction (the arena wall with visual cues is superimposed). (**G**) Logarithmic plot of fraction of turn-selective cells at different significance levels (Average ± s.e.m, 1368 cells, 4 mice, 4 FOVs). (**H**) Distribution of turn-selectivity index for all turn-selective cells (Average ± s.e.m, for p < 0.05). (**I**) Average activity of all turn-selective cells separated by CW and CCW rotations. (**J**) Example FOV showing the spatial distribution of CW (blue) and CCW-selective cells (red, for p < 0.05).

To perform large-scale recordings from neurons with high spatial resolution and without bias towards active neurons or specific cell types, which is a disadvantage of extracellular recordings, we rotated mice expressing the Ca^2+^ indicator GCaMP6s under a two-photon microscope (Figure 1D-E). We solved several technical challenges to record fluorescence transients free of image artefacts (Figure 1E, see Methods, Video S2-4). The rotation platform is ultra-stable and vibration-free such that neurons moved less than 1 µm in the focal plane during 360° rotations (see Methods). In addition, it was key to align the optical path of the custom-built two photon microscope with high precision (see Methods). Having solved these challenges, this method now enables recording the activity of neuronal somata, spines and axons of specific cell types during head rotations.

We first measured the activity of excitatory neurons in layer 2/3 of agranular RSC (1368 cells, 4 thy1-GCaMP6s mice (Dana et al., 2014)). During rotations, we found that a large fraction of neurons preferred rotations in either a CW or CCW direction (Figure 1F-G, CW = 12 %, CCW = 12 %, for p ≤ 0.05). To quantify the direction preference, we calculated a turn-selectivity index (Figure 1H, see Methods), where a value of −1 or +1 indicates that a cell only responds during CCW or CW trials, respectively. We found that only a small percentage of cells were exclusively active only in one direction. Instead, most cells were predominantly active in one direction, but also, to a lesser degree, in the other direction (Figure 1H-I). Since we always recorded in the left hemisphere, we also considered whether direction selectivity is lateralized. However, the percentages of CW and CCW cells in the same hemisphere were similar (Figure 1G-I). Finally, because recent data from macaques indicated that turn-selective neurons in posterior parietal cortex are organized in clusters (Avila et al., 2018), we tested whether CW or CCW selective cells are spatially organized. Instead of spatial clustering, we found that CW and CCW cells in mouse RSC are rather organized in a “salt and pepper” fashion (Figure 1J, Figure S1). Altogether these data show that RSC contains a substantial fraction of turn-selective neurons that are spatially intermingled.

### Many turn-selective neurons are biased by head direction

We observed two types of turn-selective neurons; Neurons that responded during CW or CCW turns regardless of the direction that the mouse was facing (see examples in Figure 1F), versus neurons that were turn-selective, but highly biased by heading direction (Figure 2A-B). The latter response type is reminiscent of the head direction cells that are known to be present in RSC (Chen et al., 1994). Of all turn-selective neurons, the majority (68 %) was significantly biased by head direction (Figure 2C, see Methods). When we selected only those cells biased by head direction and ranked them according to their preferred direction, we observed that this neuronal population represented all 360° directions evenly (Figure 2D-E). Therefore, it is unlikely that head direction tuning was influenced by a specific visual cue or by the rewarded direction. We also verified that the rotation protocol allowed the mouse to spend an equal amount of time facing each direction, thus avoiding a bias towards a specific direction by oversampling (see Methods). In summary, these data show that a significant fraction of turn-selective cells in RSC conjunctively encode information about head direction.

**Figure 2.**
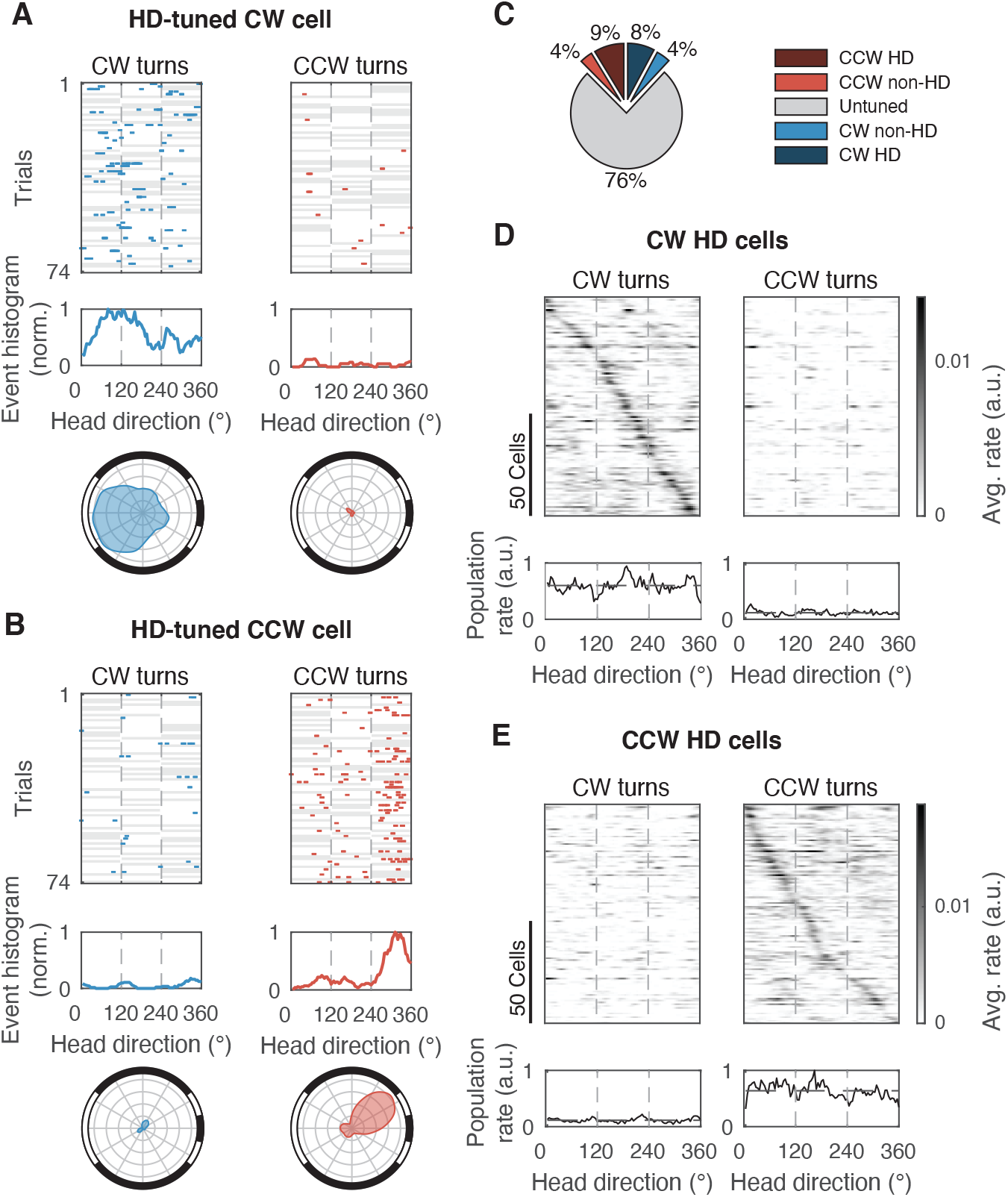
Many turn-selective neurons are biased by head direction. (**A**) Example of a CW rotation-selective cell that is biased by heading direction. Top: Deconvolved DF/F events (ticks) during CW and CCW rotation trials. White background indicates angular positions that were visited in a given trial while gray background indicates angular positions that were not visited. Middle: Event histogram for the whole session. Bottom: Polar histograms of events (Bin size = 10 degrees and smoothened using moving average over 5 bins). Dashed vertical lines indicate the three stationary positions. (**B**) Same as (A) for a CCW example cell. (**C**) Percentage of CW or CCW cells that are also biased by heading direction (HD, 1368 cells, 4 mice, 4 FOVs). (**D**) Top: Average response of all CW cells that are significantly biased by heading direction, sorted according to their preferred heading direction. Bottom: Average response of all CW cells. (E) Same for all CCW cells.

### A large fraction of turn-selective neurons encode AHV

Next, we determined whether CW and CCW-selective neurons encode not only the direction but also the angular head velocity (AHV). To reduce the influence of conjunctive head direction tuning, we limited the range of directions that the mouse could face by rotating the stage back and forth between two target positions located 180° apart (Figure 3A, 4928 cells, 10 mice). To determine the distribution of behaviourally relevant AHV, we analysed published data of freely moving mice (Laurens et al., 2019) and found that about 90 % of all head rotations are slower than 180 °/s (see Methods, n = 25 mice, see also Mallory et al. (2021)). Therefore, we rotated the mice at speeds between −180 to +180 °/s in steps of 45 °/s (Figure 3A).

**Figure 3.**
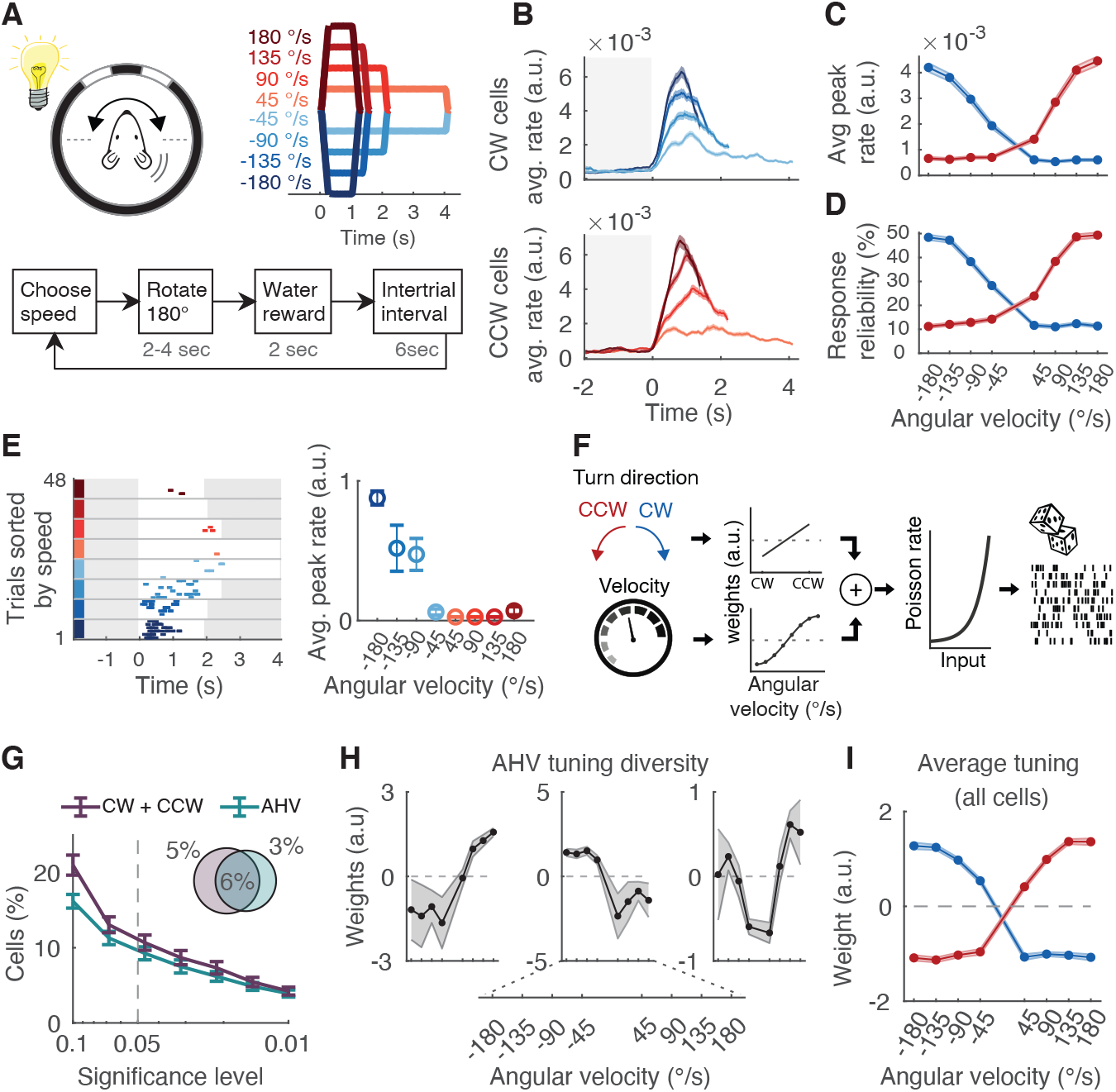
A large fraction of turn-selective neurons encode AHV. (**A**) Top: Experimental configuration; Mouse is rotated 180 degrees back and forth in CW and CCW direction, and at different rotation speeds. The speed profiles are coloured blue (CW) and red (CCW). Bottom: Block diagram of the sequence of events for a single trial. The speed is randomly chosen for every trial. (**B**) DF/F event rate (average ± s.e.m) of all cells classified as CW (top) or CCW selective (bottom), and separated by rotation speed (4928 cells, 10 mice, 14 FOVs). (**C**) Peak response (average ± s.e.m) as a function of rotation speed, for CW (blue) and CCW (red) rotations. Same data as in (B). Only the first 1 second was analysed which is the duration of the fastest rotation. (**D**) Response reliability (average ± s.e.m) as a function of rotation speed, for CW (blue) and CCW (red) rotations. Same data as in (B). (**E**) Left: Responses of an example CW cell for a whole session (DF/F events). Trials are sorted by speed. The trial starts at time = 0. White background indicates that the mouse is rotating, while a grey background indicates that the mouse is stationary. Right: Angular head velocity (AHV) tuning curve showing peak response (average ± s.e.m event rate) as a function of rotation speed. (**F**) Cartoon of Linear Non-Linear Poisson (LNP) model. The model input variables Turn direction and Velocity are converted into a weight parameter, which are summed in a linear way. Then, they are converted by a non-linear exponential into a Poisson rate and matched to the observed data (see Experimental Procedures). (**G**) Percentage of cells classified by the LNP model as turn-selective (CW and CCW cells) or tuned to AHV, as a function of significance threshold. Pie chart shows the percentage of classified cells using p < 0.05. (**H**) AHV tuning curve examples of three cells using the LNP model (average ± s.e.m). (**I**) The average ± s.e.m tuning curve of all AHV cells classified by the LNP model.

For every rotation speed, we calculated the average response of all CW and CCW-selective neurons (Figure 3B). This revealed that the response amplitude increased monotonically as a function of AHV (Figure 3C). However, the increase saturated for the highest velocities (Figure 3C). We observed a similar relationship between AHV and the response reliability (Figure 3D). In line with the average response of all CW and CCW cells, we found examples of individual neurons that had a similar monotonic relationship between response amplitude and AHV (Figure 3E).

To determine how individual cells are tuned to AHV without imposing assumptions about the shape of the neural tuning curves, we used an unbiased statistical approach (Hardcastle et al., 2017) (Figure 3F). This method fits nested linear-nonlinear-Poisson (LNP) models to the activity of each cell (see Methods). We tested the dependence of neural activity on both turn-direction (CW or CCW) and AHV (Figure 3F). From the total population, 11 % of neurons were turn-selective (Figure 3G, 527 out of 4928 cells). Among these turn-selective cells, 57 % significantly encoded AHV (303 out of 527 cells). With this model, we found that AHV tuning curves were heterogenous. Some cells had a simple monotonic relationship with angular velocity, while other cells had a more complex relationship (Figure 3H). As a sanity check of the model results, the average tuning curve of all AHV cells reproduced the simple monotonic relationship of the population response (Figure 3I, compare to Figure 3C). Altogether, these data show that a significant number of turn-selective cells encode AHV, and that while the average response of all cells has a simple monotonic relationship with AHV, the tuning curves of individual cells are heterogenous.

### AHV can be decoded from neural activity in retrosplenial cortex

We used the LNP model to quantify how well neurons in RSC report the motion stimulus. Although neurons of a single recording session could reliably discriminate CW from CCW rotations (Figure 4A, average 308 cells/session, 10 mice), the estimate of angular velocity was less reliable (Figure 4B, Video S5). We further illustrate this by comparing the actual AHV with the decoded AHV using a confusion matrix (Figure 4C). However, the reliability of reporting the correct speed was velocity dependent. Low velocity trials were more often correctly reported compared to high velocity trials (Figure 4D). For 45 °/s rotations, neurons reported the correct speed in almost 90 % of trials, while for higher speeds (90-180 °/s), this reduced to about 30-50 %. Still, these were better estimates than predicted by chance (Figure 4D). The velocity-dependent decoding of AHV was also apparent when we calculated the velocity error (Figure 4E). For low angular speeds the velocity error was about 25 °/s, while for higher speeds it increased to about 50 °/s. Another way to measure of how well a neuronal circuit reports a behavioural variable is to quantify how the decoding error scales with the population size (Figure 4F). Because we typically record 300-350 neurons simultaneously, we subsampled this population and quantified the decoding error for different population sizes. These data show that around 350 cells reported angular speeds between-180 °/s and + 180 °/s with an average error of around 30 °/s (Figure 4F). This is close to the psychophysical limit of 24 °/s that mice can discriminate in a similar task in darkness (Vélez-Fort et al., 2018). Altogether, these data indicate that a substantial amount of information about angular head movements is embedded in the RSC network, and that as few as 350 RSC neurons can provide an estimate of the actual AHV with an error of 30 °/s.

**Figure 4.**
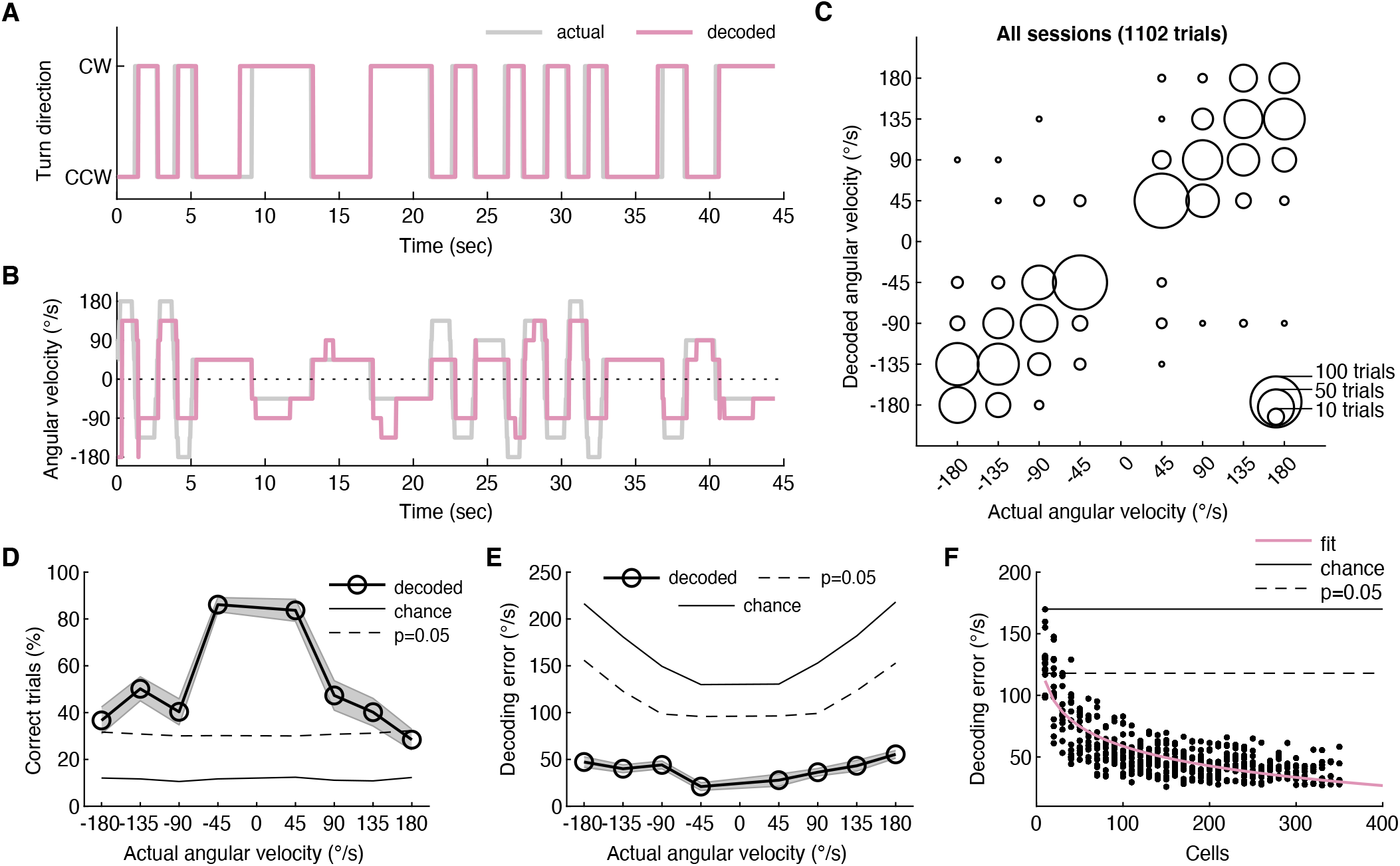
AHV can be decoded from neural activity in retrosplenial cortex. (**A**) Example trial sequence showing a mouse being rotated in the CW or CCW direction. The periods when the mouse is stationary are removed. Traces show the actual rotation direction, and that predicted by the LNP model. For decoding, all neurons in a FOV were used regardless of whether they were turn-selective or not. (**B**) Same trial sequence as shown in (A) but now showing the angular head velocity (AHV). (**C**) Scatter plot showing the actual AHV of a trial and that predicted by the LNP model. All trials of all mice are included (16 sessions, 1102 trials, 10 mice). The size of a bubble indicates the number of trials. The number of neurons used for decoding was 308 ± 15 (average ± s.e.m) per session. (**D**) The percentage of trials (average ± s.e.m) correctly decoded by the LNP model. (**E**) The decoding error (average ± s.e.m) using the LNP model as a function of actual rotation speed. (**F**) The decoding error as a function of the number of random neurons used for decoding.

### AHV tuning in retrosplenial cortex depends on sensory transduction in the vestibular organs

To determine the sensory origin of neuronal responses during rotation, we first considered whether neurons are driven by head motion or by visual flow. To distinguish between these two possibilities, we rotated mice either in complete darkness, or we kept the mice stationary and simulated visual flow by rotating the enclosure with visual cues (Figure 5A, rotation in light, 4928 cells; rotation in darkness, 4963 cells; visual flow, 3192 cells). Cells that were tuned to AHV in light, typically remained tuned to AHV in darkness, but they lost their tuning when only rotating the visual cues (Figure 5B). Similarly, when we calculated the average responses of all turn-selective neurons (281 CW cells, 246 CCW cells, n = 10 mice), we observed clear AHV tuning during rotations in both light and darkness, but not during presentation of visual flow (Figure 5C). Overall, the percentage of turn-selective and AHV cells was similar in light and darkness, but was significantly less when presenting only visual flow (Figure 5D). These data are further supported by the LNP model. We decoded AHV under all three conditions. During rotations in light or darkness, neurons reported AHV almost equally well (Figure 5E). Decoding AHV in the dark only degraded for the lowest angular velocities (45 °/s). In contrast, when we rotated only the visual cues, the decoded velocity error increased substantially, and only the lowest speeds could be decoded better than expected by chance (Figure 5E). Altogether these data suggest that AHV tuning is mostly mediated by head motion and that visual flow only contributes to low (45 °/s) rotation speeds.

**Figure 5.**
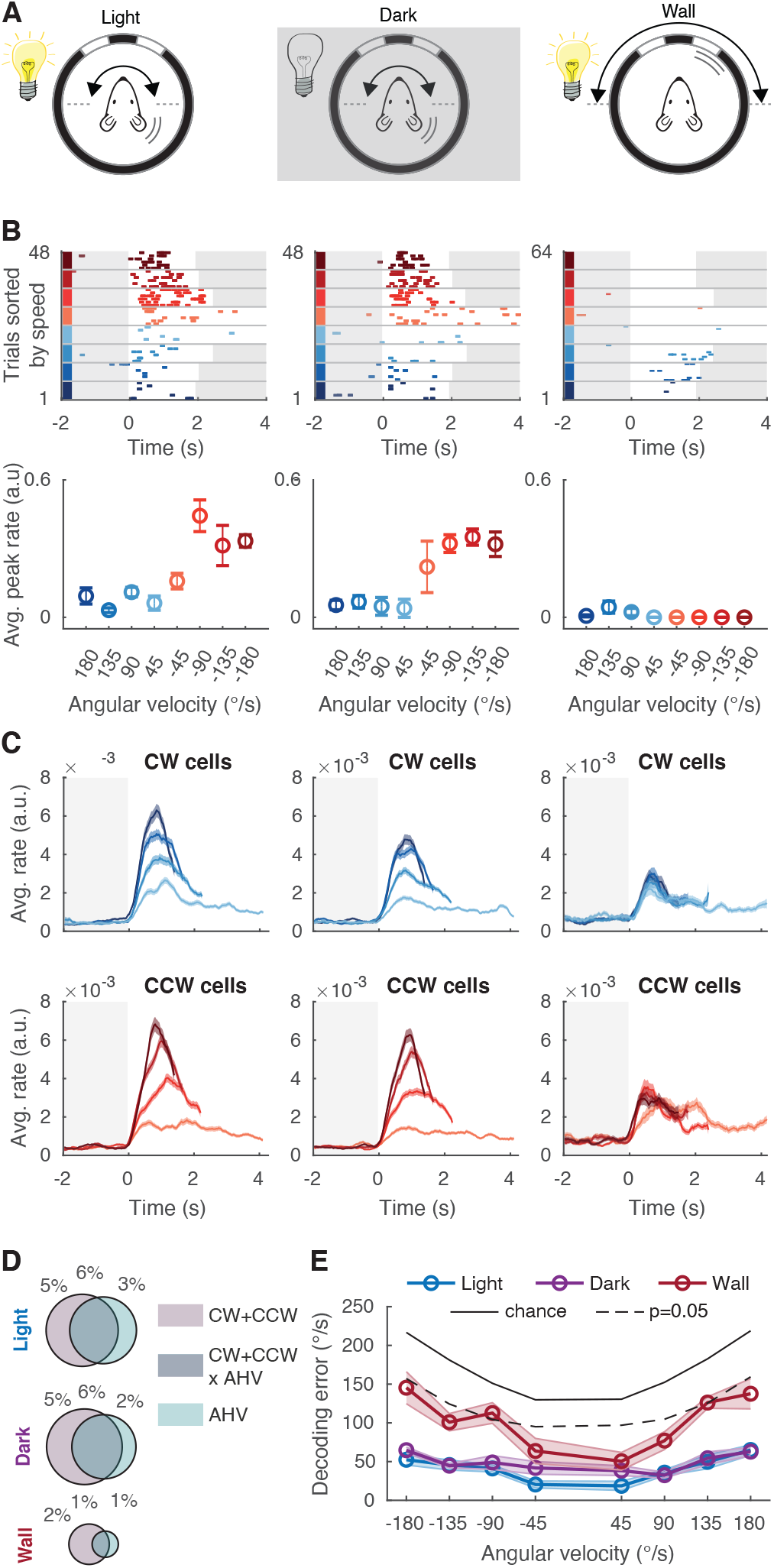
AHV tuning in retrosplenial cortex depends on sensory transduction in the vestibular organs. (**A**) Experimental configuration. Left: Mouse is rotated 180 degrees back and forth in CW and CCW direction, and at different rotation speeds. Middle: Same experiment repeated in the dark. Right: Mouse is stationary, but now the wall of the arena with visual cues is rotated to simulate the visual flow experienced when the mouse is rotating. (**B**) Top row: Responses of an example cell for a whole session (DF/F events) under the three different conditions indicated in (A). Trials are sorted by speed (see colour code in Figure 3A). White background indicates that the mouse is rotating, while a grey background indicates that the mouse is stationary. Bottom row: Angular head velocity (AHV) tuning curves showing peak response (average ± s.e.m event rate) as a function of rotation speed. (**C**) DF/F event rate (average ± s.e.m) of all cells classified as CW(top row) or CCW selective (bottom row) and separated by rotation speed. Total number recorded cells: Rotations in light = 4928 cells from 10 mice; Rotation darkness = 4963 cells from 10 mice; Visual flow-only = 3192 cells from 7 mice. (**D**) Pie charts show the average percentage of classified cells using the LNP model in each of the three conditions indicated in (A) for p < 0.05. (**E**) The AHV decoding error (average ± s.e.m) as a function of rotation speed using the LNP model.

Are neuronal responses during rotation driven by mechanical activation of the semi-circular canals in the vestibular organs? To test this, we compared neuronal responses in RSC with the responses of the vestibular nerves that convey the output of the canals to the brainstem (Figure S2A). We first simulated how mechanical activation of the canals is translated into nerve impulses in the vestibular nerve under a variety of rotation conditions (Laurens and Angelaki, 2017), and then we performed experiments to test whether neuronal responses in the RSC resemble those.

First, we rotated mice in the dark at the same speed but with different accelerations (Figure S2B). While the canals inherently sense acceleration, they transform this into a neuronal speed signal due to their biomechanical properties (Fernandez and Goldberg, 1971). Therefore, the vestibular nerve conveys speed information, at least for short lasting rotations (Figure S2C). Then, we selected only those neurons that were CW or CCW-selective and calculated their mean response (Figure S2D, 594 neurons, 4 mice). For different accelerations, the neuronal responses in RSC resembled the simulated responses of the vestibular nerve (Figure S2C), suggesting that neuronal activity in the RSC is driven by a speed signal from the vestibular organs. Next, we took advantage of the well-known observation that when the head is rotated at constant speed, the inertia signal in the canals attenuates, causing a reduction of vestibular nerve impulses (Goldberg and Fernandez, 1971) (Figure S2E, F). In mice, this occurs with a time constant of ~2.2 - 3.7 seconds (Lasker et al., 2008) (Figure S2F). To test whether RSC responses have the same temporal profile, we rotated mice at a constant speed for different durations (Figure S2G). The average response of all CW or CCW-selective neurons indeed resembled the simulated vestibular nerve activity, and decayed with a time constant of 3.1 seconds. Finally, when animals are rotated at a constant speed so that the inertial signal in the canals adapts, a sudden stop will be reported by the contralateral canals as a rotation in the opposite direction. When we analysed the average response of CW-selective neurons at the end of CCW rotations, we found indeed that a sudden stop activated these cells (Figure S2H). Altogether, these observations are fully compatible with the hypothesis that the vestibular organs drive neuronal responses in RSC during rotations in the dark.

Finally, inspired by previous work in rabbits (Sikes et al., 1988), we tested whether RSC neurons encode proprioceptive information about eye movements that occur during passive rotations (so-called "pupil position resets" as a consequence of the vestibulo-ocular reflex). Such signals may represent proprioceptive feedback, possibly mediated by stretch receptors in the eye muscles (Blumer et al., 2016). We analysed pupil-reset events during rotations in the dark using infrared illumination (Figure S3). As the rotation speed increased, we found, as expected, that the frequency of the pupil-resets also increased (Figure S3A, B). Then, to disentangle whether neuronal activity encodes head rotation or pupil resets, we performed pupil reset-triggered averaging of neuronal responses. However, this analysis provided no convincing evidence that RSC neurons report fast eye movements under our conditions (Figure S3C-F, 3708 neurons, 5 mice). In summary, these data are consistent with the hypothesis that AHV signals depend on sensory transduction in the vestibular organs, with visual flow only playing a substantial role during low velocities.

### AHV is widespread in cortical circuits

Are AHV cells restricted to RSC, or are they widespread in the cortex? To address this question, we explored large areas of the cortex with cellular resolution. Because the rotation experiments prevented us from tilting either the microscope or the mouse, we limited our exploration to the most dorsal and medial parts of the cortex (Figure 6A, 16642 cells, 15 mice). We implanted large cranial windows (~5 × 5 mm) that gave access to all imageable cortical areas (Kim et al., 2016). These included parts of primary and secondary visual cortex (V1 and V2), the posterior parts of primary and secondary motor cortex (M1 and M2), posterior parietal cortex (PPC) and the medial aspect of somatosensory cortex (S1, mostly hind limb and trunk). To align two-photon cellular-resolution images with a standardized atlas of the cortex (Kirkcaldie, 2012), we performed intrinsic optical imaging of the tail somatosensory cortex (Figure S4). This had several key advantages. Because the tail area is a small cortical area (~300 µm) closely located to the intersections of S1, M1, M2, PPC and RSC, identifying its location enables registering these areas to an atlas with high precision (Lenschow et al., 2016; Sigl-Glöckner et al., 2019) (Figure S4A-D).

**Figure 6.**
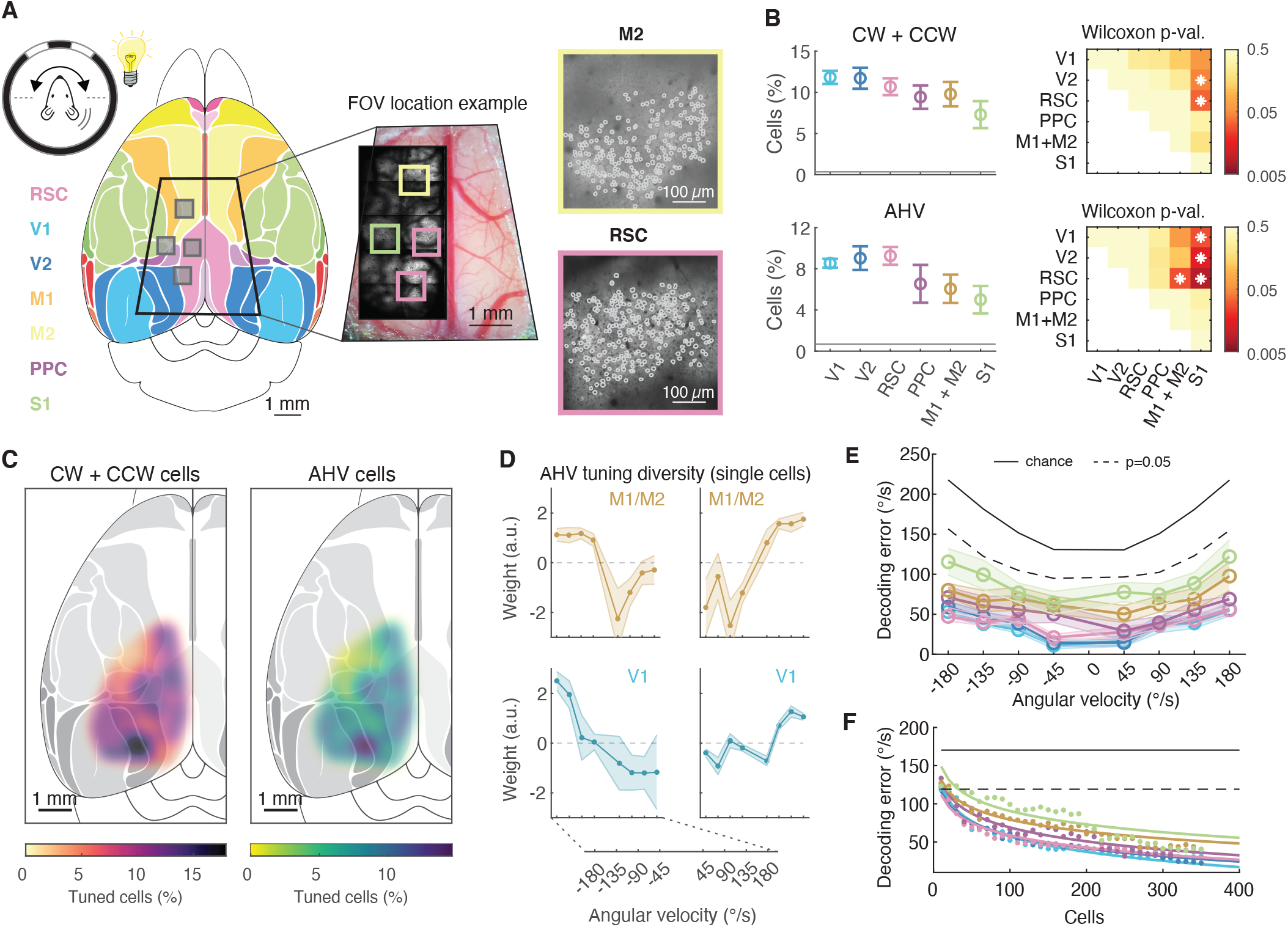
AHV is widespread in cortical circuits. (**A**) (Left) Mouse is rotated 180 degrees back and forth in CW and CCW direction, and at different rotation speeds. Map of the mouse dorsal cortex with the major cortical areas colour coded (see Kirkcaldie (2012)). Trapezoid outline indicates the size of the implanted large cranial windows. Grey boxes indicate the typical size of an imaging FOV. (Middle) Trapezoid cranial window centred on the central sinus with 4 example FOVs. (Right) Example two photon image of a FOV from secondary motor cortex (M2) and retrosplenial cortex (RSC). Cell bodies are outlined. (**B**) Left column: The percentage of turn-selective (top) and AHV-tuned cells (bottom) across different cortical areas (average ± s.e.m). The baseline indicates chance level. Right column: Statistical comparison between cortical areas (asterisk indicates pairwise comparisons with p < 0.05, Wilcoxon-Mann-Whitney test). Number of cells and mice recorded per area: V1, 2463 cells from 8 mice; V2, 4529 cells from 10 mice; RSC, 4928 cells from 10 mice; PPC, 1044 cells from 4 mice; M1+M2, 2483 cells from 7 mice; S1, 3356 cells from 9 mice. (**C**) The percentage of turn-selective (left) and AHV-tuned cells (right) mapped onto the mouse dorsal cortex. (**D**) Example AHV tuning curves obtained using the LNP model from two example areas M1+M2 and V1. (**E**) The AHV decoding error (average ± s.e.m) using the LNP model as a function of actual rotation speed, for each of the cortical areas. (**F**) The decoding error as a function of the number of random neurons used for decoding.

We rotated mice at different speeds in light conditions (as in Figure 3A), and we found neurons that reported the direction and speed of motion in all areas that we explored (Figure 6B, C). Compared to RSC, the percentage of cells in V1 and V2 was very similar (Figure 6B), but the numbers in PPC, M1/M2 and S1 were lower (Figure 6B). To quantify how well neurons in these different areas report AHV, we used the LNP model (Figure 6D-F). Neurons in all areas decoded AHV better than expected by chance (Figure 6E). However, while neurons in RSC, V1, V2 and PPC reported AHV equally well, neurons in M1/M2 and notably S1 performed worse (Figure 6E). The overall number of recorded neurons per session was slightly different between cortical areas and this can affect the decoding performance. To account for this, we down-sampled the number of neurons in individual recording sessions and quantified the decoding error as a function of population size (Figure 6F). These data show that, with the exception of M1/M2 and S1, a population size of 300-350 neurons in each area can report AHV with a performance close to the psychophysical limits for discriminating angular speed in mice (Vélez-Fort et al., 2018). Thus, altogether, angular direction tuning and AHV are widespread in cortical circuits, but with some differences in how well AHV can be decoded from the neural population.

### The sensory origin of AHV tuning is cortical area-dependent

To test whether AHV tuning in different cortical areas depends on head motion or visual flow, we compared neuronal responses when rotating mice in light or darkness, or when rotating only the visual cues (Figure 7A). Maps of AHV-tuned cells in these three conditions showed notable differences between brain areas (Figure 7B). Compared to rotations in light, rotations in the dark reduced the percentage of AHV cells in V1 and V2 but did not substantially affect other brain areas (Figure 7C, D). Conversely, when rotating only the visual cues, the percentage of AHV cells in V1 remained the same, strongly reduced in V2, and was near-zero in all other brain areas. These data show that with the exception of V1 and V2 where AHV tuning is strongly dependent on visual flow, AHV tuning in all other areas (PPC, RSC, M1/M2 and S1) depends mostly on head motion.

**Figure 7.**
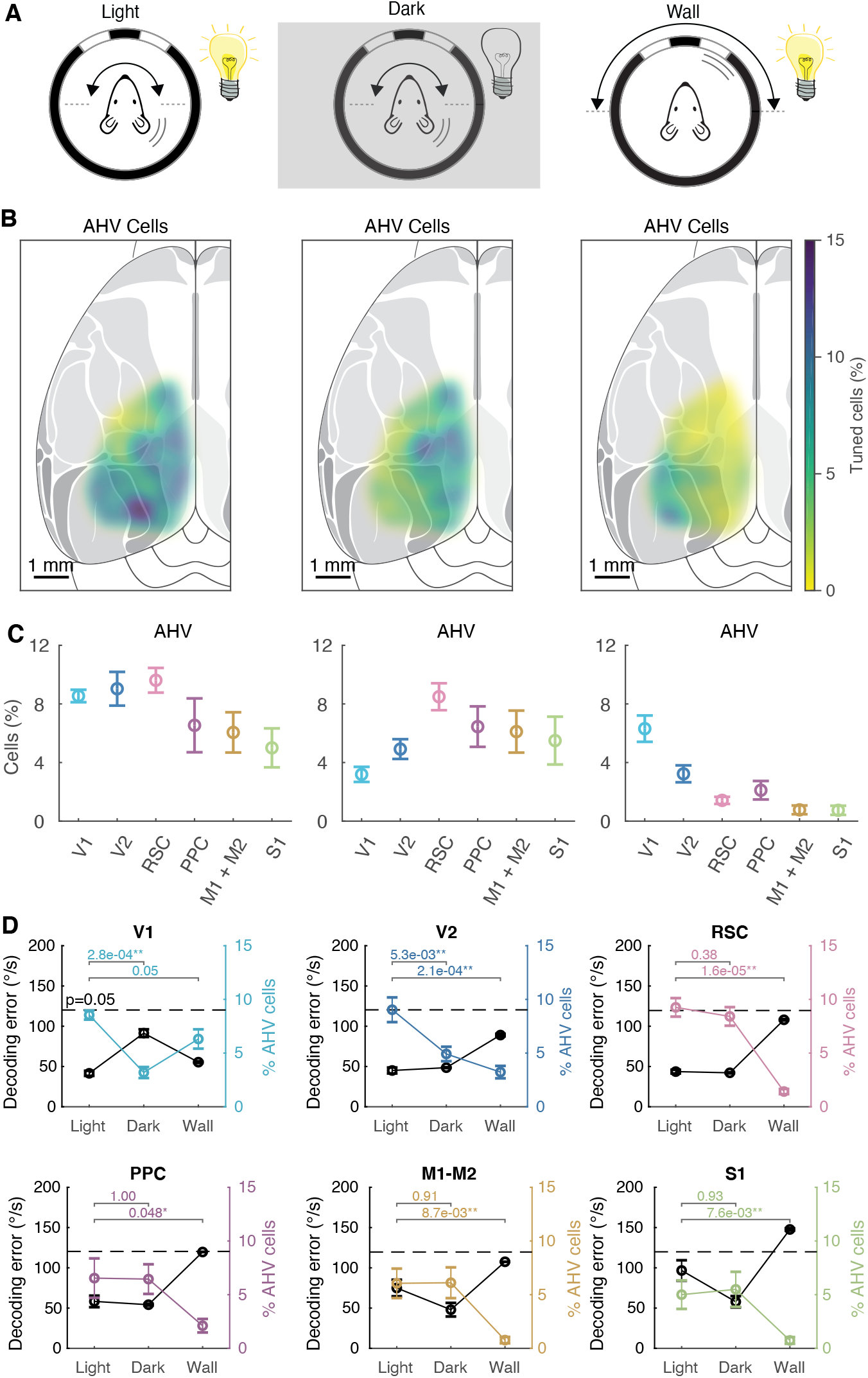
Whether AHV tuning depends on head motion or visual flow is area dependent. (**A**) Experimental configuration. Left: Mouse is rotated 180 degrees back and forth in CW and CCW direction, and at different rotation speeds. Middle: Same experiment repeated in the dark. Right: Mouse is stationary, but now the wall of the arena with visual cues is rotated to simulate the visual flow experienced when the mouse is rotating. (**B**) The percentage of AHV-tuned neurons mapped onto the mouse dorsal cortex, tested in each of the three conditions. (**C**) The percentage of AHV-tuned cells across different cortical areas (average ± s.e.m) tested in each of the three conditions. (**D**) The decoding error (black, average ± s.e.m) and percentage of AHV cells (colour coded) under the three different conditions. The horizontal bars above show the Mann-Whitney unpaired test p-values for comparison of percentage of AHV cells across different conditions. The dashed lines show the p=0.05 level for decoding errors. The number of cells and mice recorded under light conditions is given in the legend of Figure 6. Under dark conditions: V1, 2385 cells from 8 mice; V2,4564 cells from 10 mice; RSC, 4963 cells from 10 mice; PPC, 1044 cells from 4 mice; M1+M2, 2489 cells from 7 mice; S1, 3398 cells from 9 mice. During wall rotations: V1, 1864 cells from 6 mice; V2, 3560 cells from 8 mice; RSC, 3192 cells from 7 mice; PPC, 816 cells from 3 mice;M1+M2, 1095 cells from 4 mice; S1, 1562 cells from mice.

Finally, we used the LNP model to test whether AHV decoding in these three different conditions varied between brain regions (Figure 7D). With the exception of V1 and V2, rotations in the dark did not affect the decoding error in any of the brain areas. In contrast, rotating only the visual cues increased the decoding error in all cortical areas, except area V1. In summary, whereas AHV tuning in V1, and to a lesser degree V2, depends mostly on visual flow, in all other cortical areas that we explored (PPC, RSC, M1/M2 and S1) AHV tuning depends mostly on head motion.

## Discussion

Here, we developed a method to measure AHV signals in the cortex using two-photon microscopy. We show that cells in layer 2/3 of agranular RSC report the direction and speed of angular head movements. Many of these cells were also influenced by the direction that the mouse was facing. Furthermore, approximately 300-350 RSC neurons were sufficient to report AHV with a precision close to psychophysical limits. AHV coding was not restricted to RSC but was present in layer 2/3 of all cortical areas explored, including substantial parts of visual (V1/V2), posterior-parietal (PPC), somatosensory (S1) and motor cortices (M1/M2). There were, however, notable differences between these areas. While AHV cells in RSC, PPC, M1/M2 and S1 depended mainly on head motion driven by the vestibular organs, AHV cells in V1 and V2 depended on visual flow.

Cortical AHV neurons that track the activity of the vestibular organs were first reported in head-fixed monkeys and cats decades ago (Büttner and Buettner, 1978; Vanni-Mercier and Magnin, 1982). Later, AHV neurons were also found in the rodent cortex, specifically in preand postsubiculum (Preston-Ferrer et al., 2016; Sharp, 1996), entorhinal cortex (Mallory et al., 2021), posterior parietal cortex (Wilber et al., 2014), anterior cingulate cortex (Mehlman et al., 2019) and RSC (Cho and Sharp, 2001). However, the underlying modalities that drive tuning have remained unknown. This was recently investigated by studying motion-sensitive neurons in area V1 of head-fixed mice (Bouvier et al., 2020; Vélez-Fort et al., 2018). By rotating mice in the dark, they show that neurons in deep cortical layers 5 and 6 were excited, while neurons in superficial layers 2/3 were suppressed. Consistent with these data, we found that fewer neurons in layer 2/3 of V1 encode AHV in the dark. However, we did not observe neuronal suppressions. Given the lo basal firing rates of neurons in superficial layers, and hence the large number of repetitions necessary to observe a suppressive effect, it is possible that suppressions remained undetected. Altogether, our work extends current knowledge by showing that cortical AHV cells are far more widespread than previously reported, spanning visual, somatosensory and motor areas, and rely on vestibular or visual input in an area-dependent manner.

By comparing rotations in light and darkness, we show that vestibular input contributes most of the information to decode AHV from neurons in RSC (Figure 5E). However, this may depend on the range of AHV that we explored. In-deed, visual flow selectively improves decoding during lower AHV (45 °/s, Figure 5E). This could indicate that vestibular and visual information play complementary roles to support a wider range of AHV. This is consistent with classic work studying compensatory eye movements as a read-out of AHV in sub-cortical circuits (Stahl, 2004). These data show that, during the vestibulo-ocular reflex, the vestibular organs report slow head rotations poorly, while during the optokinetic reflex, sub-cortical circuits report fast visual flow poorly (> 40 °/s). Therefore, AHV tuning driven by different modalities may not be redundant but play complementary roles by expanding the bandwidth of AHV representations.

We found that AHV cells in many cortical areas depend on vestibular input. But how can vestibular signals be so widely distributed? Recent work suggested that vestibular-dependent AHV cells in V1 are driven by neurons in RSC, which in turn receive AHV input from the anterodorsal thalamic nucleus (ADN) (Vélez-Fort et al., 2018). It is generally assumed that ADN encodes angular head motion because it receives input from the lateral mammillary nucleus, which contains many AHV cells (Cullen and Taube, 2017; Stackman and Taube, 1998). However, while ADN has been extensively studied, AHV cells in ADN have not been reported. Therefore, it is unclear how AHV could be transmitted via this route. An alternative pathway is the more direct system of vestibulo-thalamocortical or the cerebellothalamocortical projections (Lopez et al., 2011). Neurons in the vestibular nucleus sent projections to various thalamic nuclei that target numerous cortical areas (Nagata, 1986; Shiroyama et al., 1999). This broad projection pattern may explain why AHV cells are so widely distributed in the cortex. However, these vestibular pathways remain poorly understood and need to be functionally characterized to determine which convey AHV. This could be achieved in a projection-specific manner by imaging thalamocortical axons in the cortex using two-photon microscopy as described here.

Animals may use cortical AHV signals to determine whether a changing visual stimulus is due to their own movement or due to a moving object. The brain likely implements this by building internal models that, based on experience, compare intended motor commands (efference copies) with the sensory consequences of actual movements (Holst and Mittelstaedt, 1950; Miall and Wolpert, 1996). Such efference copies could then anticipate and cancel self-generated sensory input that would otherwise be unnecessarily salient. Consistent with this hypothesis, a classic example is the response of vestibular neurons in the brainstem that, compared to passive rotation, are attenuated during voluntary head movements (Brooks et al., 2015; Medrea and Cullen, 2013; Roy and Cullen, 2001). This raises questions about the role of AHV signals recorded during passive rotation. However, internal models also predict that, during movements, the ever-ongoing comparison between expected and actual sensory stimuli often results in a mismatch. A necessary corollary is that AHV signals during passive rotation are essentially mismatch signals because the lack of intended movements cannot cancel the incoming vestibular input. Therefore, vestibular mismatch signals may play two fundamental roles: To update self-motion estimates and to correct motor commands, and to provide a teaching signal to calibrate internal models for motor control and so facilitate motor learning (Miall and Wolpert, 1996). Both these functions likely require that cortical circuits have access to AHV information.

The widespread existence of AHV cells in the cortex also has implications for how animals orient while navigating through an environment. Attractor network-dependent head direction models make use of neurons that convey the direction and speed of angular motion (Hulse and Jayaraman, 2019; Knierim and Zhang, 2012; Taube, 2007). These neu-rons dynamically update head direction cell activity in accordance with the animal’s movement in space. In mammals, the mainstream hypothesis positions the attractor network in circuits spanning the mesencephalon and hypothalamus, in part because that is where AHV cells are commonly found (Cullen and Taube, 2017; Taube, 2007). Indeed, the limited and often anecdotal evidence of AHV cells in the rodent cortex has led to the widespread notion that angular velocity signals are largely restricted to sub-cortical areas (Cullen and Taube, 2017; Hulse and Jayaraman, 2019). However, several studies report cortical neurons encoding turn direction, but whether these cells also encode velocity is not known (Alexander and Nitz, 2015; McNaughton et al., 1994; Whitlock et al., 2012). In addition to AHV cells, attractor networks also need cells that conjunctively code AHV and head direction (Hulse and Jayaraman, 2019; McNaughton et al., 1991; Taube, 2007). Such cells were previously discovered in subcortical structures such as the lateral mammillary nucleus (Stackman and Taube, 1998) and dorsal tegmental nucleus (Bassett and Taube, 2001; Sharp et al., 2001). There is, however, also evidence for such cells in the cortex (Angelaki et al., 2020; Cho and Sharp, 2001; Sharp, 1996; Wilber et al., 2014). Here, we show that AHV cells are far more common in cortical circuits than previously thought. We also report a considerable fraction of cells in RSC that conjunctively encode turn-selectivity and head direction. Altogether, these data indicate that the basic components of the attractor network architecture are not limited to subcortical structures but are also present in the cortex. Therefore, our findings add to the debate about which circuits are involved in generating head direction tuning.

In conclusion, our results show that cortical AHV cells are far more widespread than previously thought, spanning a large fraction of cortex, and rely on vestibular or visual input in an area-dependent manner. How the distributed nature of AHV cells contribute to calibrating internal models for motor control and movement perception and generating head direction signals are important questions that remain to be addressed. The ability to perform two-photon imaging in head-fixed animals subjected to rotations and in freely moving animals (Voigts and Harnett, 2019; Zong et al., 2017), will provide exciting new opportunities to explore these important questions at both the population and sub-cellular level in a cell type-dependent manner.

## ACKNOWLEDGEMENTS

This work was funded by the European Research Council (ERC Starting Grant #639272 to K.V., the Research Council of Norway #231495, #276047 and #274306 to K.V.). We thank Kristin Larsen Sand, Rune Lanton and Michele Gianatti for technical assistance. We thank Bruno Pichler (Independent NeuroScience Services; INSS) for developing the custom two-photon microscope and providing technical assistance. Some schematics were created with Biorender.com. Finally, we thank Tsai-Wen Chen, Lyle Graham, Jean Laurens, Nuo Li, Raul Muresan and Ninglong Xu for providing critical comments that significantly improved a draft of the manuscript.

## AUTHOR CONTRIBUTIONS

E.H, A.W, A.C and K.V designed research. E.H. performed all experiments and model-free data analysis. A.W. performed all model-based data analysis. A.C. performed pilot experiments and data analysis. K.V. performed surgeries. E.H., A.W. and K.V. wrote the paper.

## Methods

### Resource availability

The data and code generated in this study are available at [name of repository] and Github, respectively [accession code / web link]. Technical drawings of the setup and microscope (Autodesk Inventor / Zemax) are available at [name of repository].

### Animals

Data includes recordings from 18 Thy1-GCaMP6s mice (Dana et al., 2014) (16 male, 2 female, Jackson Laboratory, GP4.3 mouse line #024275). Mice were 2-5 months old at the time of surgery and were always housed with littermates in groups of 2-5. Mice were kept on a reversed 12hour light/ 12-hour dark cycle, and experiments were carried out during their dark phase. To keep mice engaged during experiments, they received water drop rewards throughout a session and therefore, they were kept on a water restriction regime as described in Guo et al. (2014). All procedures were approved by the Norwegian Food Safety Authority (projects #FOTS 6590, 7480, 19129). Experiments were performed in accordance with the Norwegian Animal Welfare Act.

## Hardware and data acquisition

### Rotation arena

#### The rotation platform

(Methods Figure 1) consisted of a circular aluminum breadboard (300 mm diameter, Thorlabs, MBR300/M) fixed on a high-load rotary stage (Steinmeyer Mechatronik, DT240-TM). The stage was driven by a direct-drive motor. This has the important benefit that the motor does not have a mechanical transmission of force and hence is vibration-free, which is critical during imaging. We placed the mice in a head post and clamp system (Guo et al., 2014) on a two-axis micro-adjustable linear translation stage (Thorlabs, XYR1), which was mounted on the rotation platform in a position where the head of the mouse was close to the center of the platform and the axis of rotation. The linear stage was necessary to micro-adjust the position of the mouse so that the field of view (FOV) for imaging was perfectly centered on the axis of rotation.

To verify that the sample stays in the focal plane during rotations, we tested the stability and the precision of the rotation platform by imaging a microscope slide with pollen grains (~30 μm diameter). We rotated the slide from 0 degrees to 360 degrees in 45-degree steps, and in each orientation, we took a z-stack of images. We extracted the fluorescence profile along the z-axis for each pollen grain, and fitted each profile with a Gaussian function to determine the center position of the pollen along the z-axis (We assumed that the center position corresponded with the location of the peak in the Gaussian profile). Finally, by determining the variance of the center positions along the z-axis for the different angular positions, we concluded that each pollen moved less than 1 μm along the z-axis during 360-degree rotations. A sub-μm quantification was prohibited by the limited precision of the z-axis stepper motors of the microscope, which was 1 μm.

#### Commutator

Electrical signals for devices mounted on the rotation platform were carried through a commutator (MUSB2122-S10, MOFLON Technologies) to allow continuous rotations without cable twisting. The commutator had 10 analog and 2 USB channels and was placed in a hollow cavity within the rotary stage. This configuration provided unlimited rotational freedom in both directions.

#### Pupil tracking camera

An infrared-sensitive camera (Basler dart 1600-60um) was mounted on the rotation platform. To avoid that the camera occupied the mouse’s field of vision, we positioned it behind the mouse, and the pupil was recorded through a so-called hot mirror (#43-956, Edmund Optics) in front of the mouse. This mirror is transparent in the visible range but reflects infrared light. The frame rate was 30 fps, and data were acquired using custom-written LabVIEW code (National Instruments). During post-processing, we measured the pupil position and diameter using scripts in MATLAB. The pupil was back-illuminated by the infrared light from the two-photon laser during imaging experiments, which created a high contrast image of the pupil.

#### Mouse body camera

To track the overall movements of the mouse, we used an infrared camera (25 fps) placed behind the mouse. Body movement was quantified in MATLAB by calculating the absolute sum of the frame-by-frame difference in pixel values within a region of interest centered on the mouse’s back.

#### Lick port

To keep the mouse engaged and keep arousal levels steady during experiments, the mouse received water rewards through a lick spout. Water delivery was controlled by a solenoid valve. The lick spout was part of a custom-built electrical circuit that detected when the tongue contacted the lick spout (Guo et al., 2014).

#### Arena wall and roof

The arena was enclosed by a 10 cm high cylindrical wall (30 cm diameter) with visual cues. The wall was custom-built and consisted of a black 2 mm thick plastic plate wrapped around and fixed to a circular "Lazy-Susan" ball bearing (30 cm diameter). The visual cues where 7 cm and 20 cm-wide white plastic cards. The height of the visual cues was restricted to avoid triggering an optokinetic reflex when moving the walls while the mouse is stationary. The walls were driven by a stepper motor (SMD-7611, National Instruments), and a rotary encoder (National Instruments) was used to read the wall position.

An illuminated roof covered the arena to prevent the mouse from observing cues outside the arena. The roof was made from two snug-fitting pieces of frosted plexiglass with an array of white LEDs mounted on top. This provided an evenly dim illumination of the arena without creating fixed visual cues. The roof was resting on top of the walls and therefore rotated together with the walls.

#### Light-shield

Because the arena was illuminated, we designed a light shield to protect the light-sensitive photomultipliers (PMTs) of the microscope. The light-shield consisted of a 3D-printed cone-shaped cup that clamped on the titanium head bar with neodymium magnets and fitted snugly over the microscope objective.

### Two-photon microscope

We used a custom-built two-photon microscope, designed with two key objectives: 1) to provide enough space around the objective to accommodate the large rotation arena and 2) provide easy access for aligning the laser beam through the optics of the excitation path.

**Methods Figure 1.**
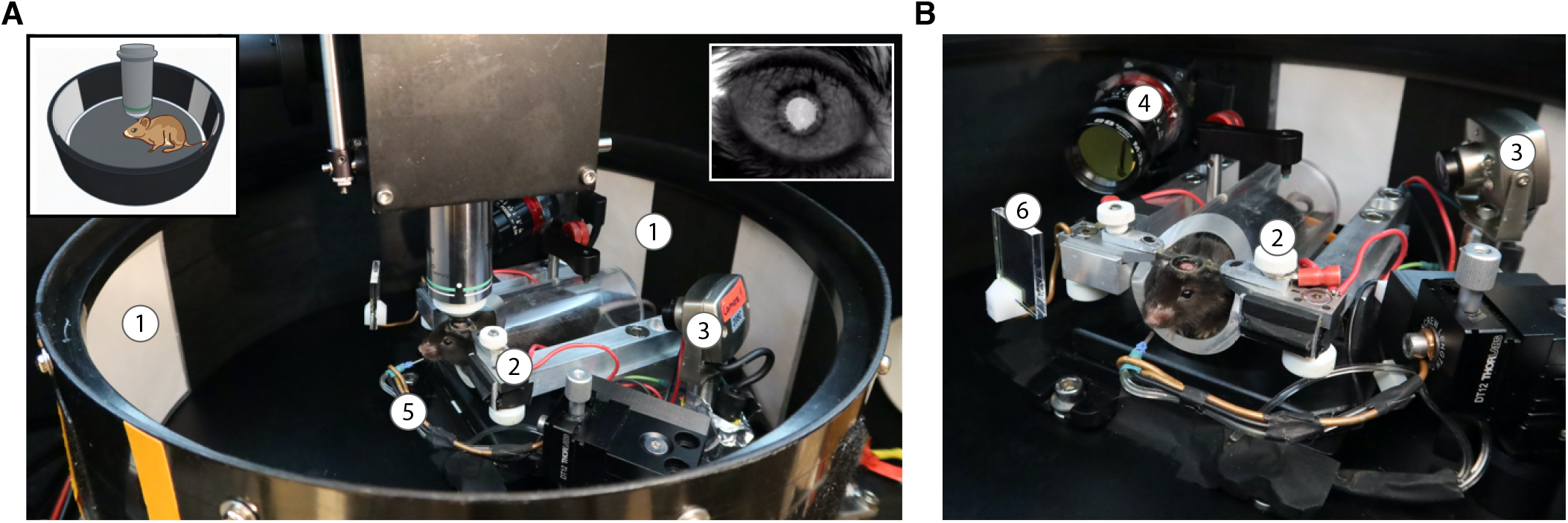
Photograph of the setup. (**A**) Overview of the mouse in the rotation arena. (**B**) Close-up of the mouse. 1 – Visual Cues, 2 – Mouse fixation frame, 3 – Body camera, 4 – Pupil camera, 5 – Lick spout, 6 – Infrared “hot” mirror.

The latter was critical for reducing fluorescence artefacts in the recording that arose from rotating the sample. The microscope was built in collaboration with Independent Neuroscience Services (INSS). The microscope consists of Thorlabs components and a parts list, Autodesk Inventor files, and Zemax files of the optical design are available [accession code/web link].

Rotating a sample while imaging can create two types of recording artefacts. The most intuitive, and the easiest to fix, is an asymmetry in the sample illumination. Due to vignetting, the illumination of the sample is usually not uniform, but follows a 2D Gaussian profile. Suppose the point of maximum brightness is offset from the image center. In that case, parts of the sample will become brighter and dimmer during rotation, giving rise to an artefact in the fluorescence signal. Another, less intuitive artefact, arises if one considers the angle of the laser beam when parked in its center position. If this angle is not perfectly parallel with the axis of rotation, then, even deviations of 1 degree will give rise to a severe luminance artefact. While by itself not critical, it is the combination of this angle offset and inhomogeneous brain tissue that leads to this type of artefact. An example of this artefact is shown in Suppl. Video S3, and an intuitive explanation is shown in Suppl. Video S4. While it is generally assumed that the laser beam is perfectly centered through all the optics of the excitation path, in practice this is rarely the case, at least not with the precision required to scan a rotating sample. Finally, it is important to note that the laser beam’s precise alignment through the excitation path optics requires easy access, which can be difficult with commercial microscopes. Hence, our motivation to design a custom microscope.

### Control and acquisition software

The rotary stage and all associated components were controlled by a custom-written LabVIEW program (LabVIEW 2013, NI). Except for the rotary stage, which had its own controller, control signals for other components were sent through a DAQ (X-Series, PCIe 6351, NI). The two-photon frame-clock, lick-port signals and wall positions were all acquired using the DAQ. The twophoton frame clock signal was used to synchronize the twophoton recording with the recorded signals from the DAQ. The angular position of the rotation stage was recorded independently through the software for two-photon acquisition.

## Experimental protocol

### Surgery

Surgery was carried out under isoflurane anaesthesia (3 % induction, 1 % maintenance), while body temperature was maintained at 37 °C with a heating pad (Harvard Apparatus). Subcutaneous injection of 0.1 mL Marcaine (bupivacaine 0.25 % *m/V* in sterile water) was delivered at scalp incision sites. Post-operative injections of analgesia (Temgesic, 0.1 mg/kg) were administered subcutaneously.

Several weeks before imaging, mice underwent surgery to receive a head bar and cranial window implant. The cranial window was either a 2.5 mm diameter round glass (7 mice) or a 5×5 mm trapezoidal glass (11 mice, see Figures 6 and 7). The center of the round window was −2.2 mm AP, and the front edge of the trapezoid window was +1.0 AP. Both windows consisted of two pieces of custom-cut #1.5 coverslip glass (each ~170 μm thick) glued together with optical adhesive (Norland Optical Adhesive, ThorLabs NOA61). The aim was to create a plug-like implant such that the inner piece of glass fits in the craniotomy, flush with the surface of the brain, while the outer piece rests on the skull. Therefore, the outer piece of the window extended by 0.5 mm compared to the inner window. Heated agar (1 %; Sigma #A6877) was applied to seal any open spaces between the skull and edges of the window, and the window was fixed to the skull with cyanoacrylate glue. A detailed protocol of the surgery can be found in (Holtmaat et al., 2009).

### Water restriction

Starting minimum 1 week after surgery, mice were placed on water restriction (described in Guo et al. (2014)). Mice were given 1-1.5 mL water once per day while body weight was monitored and maintained at 10-15 % of the initial body weight. During experiments, mice would receive water drops of 5 μl to keep them engaged.

### Animal training / habituation

During the first week of water restriction, mice were picked up and handled daily to make them comfortable with the person performing the experiments. Before experiments started, mice were placed in the arena and allowed to explore it for about 5 minutes. Next, they were head-fixed for about 30 minutes to screen the window implant. Experimental sessions would then start the next day.

### Two-photon imaging

All imaging data were acquired at 31 fps (512×512 pixels) using a custom-built two-photon microscope and acquisition software called SciScan (opensource, written in LabVIEW). The excitation wavelength was 950 nm using a MaiTai DeepSee ePH DS laser (SpectraPhysics). The average power as measured under the objective (N16XLWD-PF, Nikon) was typically 50-100 mW. Photons were detected using GaAsP photomultiplier tubes (PMT2101/M, Thorlabs). The primary dichroic mirror was a 700 nm LP (Chroma), and the photon detection path consisted of a 680 nm SP filter (Chroma), a 565 nm LP dichroic mirror (Chroma), and a 510/80 BP filter (Chroma). The FOV size varied between 500-700 μm.

All data were recorded 100-170 μm below the cortical surface, corresponding to L2/3 of agranular RSC. At the start of a recording, we adjusted the mouse position to align the FOV center with the axis of rotation. This required three steps: 1) Before placing the mouse, we scanned a rotating slide with 30 μm pollen grains to align the axis of the objective with the center of rotation of the FOV. 2) Because positioning the mouse caused small (< 100 μm) displacements, a final fine adjustment of the objective position is needed. This was done automatically by recording an average projection image stack at 0° and 180°, respectively, and then aligning the de-rotated images to estimate the offset from center. 3) When the objective was in place, we manually adjusted the position of the mouse (using the translation platform) to find a FOV and avoiding areas with large blood vessels.

### Allocentric rotation protocol

Data presented in Figures 1-2 were recorded during 360-degree rotations. Mice were rotated in randomly selected steps of 120, 240 or 360 degrees within a range of ±720 degrees. The angular rotation speed was 45 °/s, and the acceleration was 200 °/s2. This ensured that most of the rotation was taking place at a constant speed. Mice were stationary for 4-8 seconds between rotations while facing 1 of 3 possible directions (0, 120 and 240 degrees). The mouse received a 5 μL water reward if its tongue touched the lick spout while stationary in the 0-degree direction. Mice typically performed 120-150 trials before they lost interest in water and the experiment was terminated.

### Angular velocity protocol

Data presented in Figures 3-7 were recorded while rotating the mice back and forth in 180degree steps (alternating CW and CCW turns) at four different speeds. The speed for each turn was selected randomly from 45 °/s to 180 °/s in steps of 45 °/s. We chose this range of speeds based on data from freely moving mice (Methods Figure 2). Angular head acceleration varied between 300-800

**Methods Figure 2.**
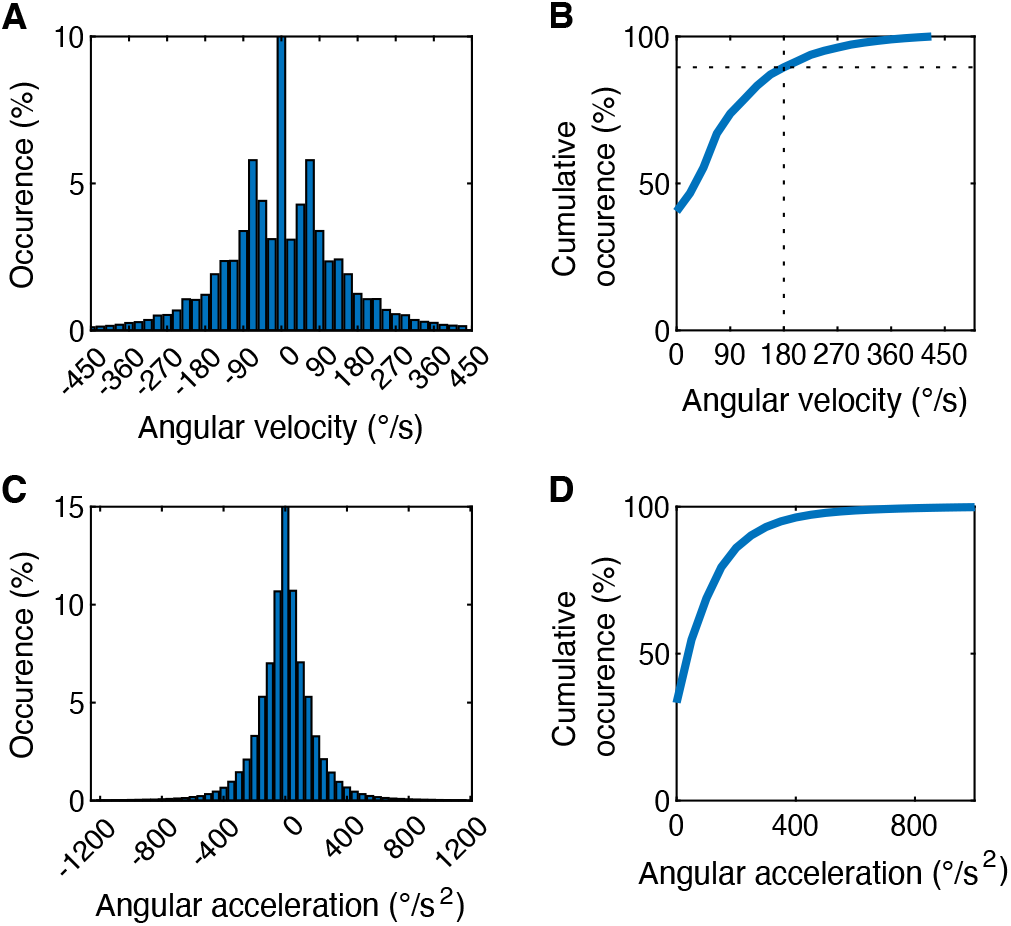
AHV and acceleration recorded from freely moving mice. (**A**) Histogram of AHV (**B**) Cumulative histogram shows that 90 % of angular movements are below 180 °/s. (**C**) Histogram of angular head acceleration. (**D**) Cumulative histogram of angular head accelerations. Data kindly provided by Jean Laurens.

°/s2 depending on which angular velocity was selected. During higher speeds, the acceleration was increased to ensure that a longer rotation period took place at a constant speed. Rotations were interleaved with 8-10 s breaks during which mice received a 5 μL water drop.

In order to separate the contributions of vestibular and visual inputs in angular head velocity tuning, each session consisted of three consecutive blocks of 50-64 trials. First, the mouse was passively rotated in light conditions, then the mouse was passively rotated in darkness and finally the walls were rotated while the mouse was stationary. For the wall rotation block, the wall was rotated using the same motion profile that was used with the rotary stage.

During the early explorations, we observed that the number of turn-selective cells consistently decreased during the first three days of recording but then remained stable (Methods Figure 3). This may be related to the mice’s overall arousal levels as they become more familiar with the apparatus. To prevent his from influencing the percentage of cells measured in different brain regions (which required recording across several days in the same mouse), we did not include sessions recorded in the first three days for this analysis.

### Registration of two-photon FOVs across brain regions

The FOV positions were registered onto a map of the dorsal brain surface in two steps: 1) A picture of the blood vessels in the cranial window was anchored to the map based on stereotaxic coordinates determined during surgery. 2) Next, twophoton images tiling the brain surface were also positioned on the map by matching the vessel pattern (see example in Figure 6A, middle panel). Finally, we used intrinsic imaging to verify the accuracy of this approach.

**Methods Figure 3.**
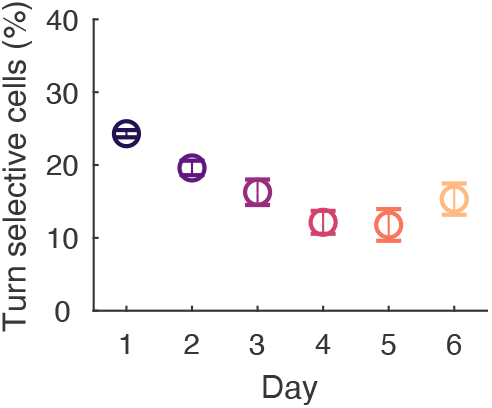
Percentage of turn-selective cells decreases over days. Mean ± s.e.m. percentage of turn selective cells observed across 6 days (n = 4 mice). Cells were classified using the LNP Model at a significance level of p < 0.05.

### Intrinsic imaging

To verify the accuracy of the FOV registration, we performed intrinsic imaging of the tail somatosensory cortex in a subset of mice (6 mice, see Figure S4). Mice were injected with the sedative (chlorprothixene, 1 mg/kg intramuscular) and maintained on a low concentration of isoflurane (~0.5 %). The mice rested on a heating pad, and their eyes were covered with Vaseline. Images were acquired through the cranial window using a CCD camera (Hamamatsu, ImagEM X2) mounted on a Leica MZ12.5 stereo microscope. Intrinsic signals were obtained using 630 nm red LED light, while images of the blood vessels were obtained using 510 nm green LED light. We stimulated the tail with a cotton swap and recorded the intrinsic signal over 5 repetitions. Each repetition consisted of a 10-second baseline recording, a 15-second stimulation period and a 20-second pause. The final intrinsic signal image was the difference between the average projection of all stimulation images and the average projection of all baseline images. Using this method, the tail somatosensory cortex obtained by intrinsic imaging was within 100-200 μm of the expected tail region based on stereotactic coordinates.

### Data processing and analysis

#### Image registration

To correct image distortion due to rotation of the sample and brain movements in awake mice, the images were registered using a combination of custom written and published code (Pnevmatikakis and Giovannucci, 2017) following three steps. 1) *Image de-stretching*: The resonance scan mirror follows a sinusoidal speed profile which distorts the images along the x-axis. This was corrected using a lookup table generated from scanning a microgrid. 2) *Image de-rotation*: During post-processing, images were first de-rotated using the angular positions recorded from the rotary stage. Additional jitter in the angular position was corrected for using FFT-based image registration (NoRMCorre, (Pnevmatikakis and Giovannucci, 2017)). Finally, to correct rotational stretch of images due to the rotation speed, a custom MATLAB script was used for a line-by-line correction. 3) *Motion correction*: De-rotated images were motioncorrected using a combination of custom-written scripts and NoRMCorre (Pnevmatikakis and Giovannucci, 2017).

#### Image segmentation

Registered images were segmented using custom-written MATLAB scripts. Soma regions of interest (ROI) were either drawn manually or detected automatically. All ROIs were inspected visually. When ROIs were overlapping, the overlapping parts were excluded. A doughnut-shaped ROI of surrounding neuropil was automatically created for each soma ROI, by dilating the soma ROI so that the doughnut area was four times larger than the soma ROI area. If this doughnut overlapped with another soma ROI, then that soma ROI was excluded from the doughnut.

#### Signal and event extraction

ROI fluorescence signals were calculated by averaging the pixel values within the ROIs for each imaging frame. ROI fluorescence changes were defined as a fractional change *DF/F* (*t*) = (*F* (*t*) − *F*_0_)*/F*_0_, with *F*_0_ being the baseline defined as the 20th percentile of the ROI fluorescence signal, *F* (*t*). This was calculated for both the soma and neuropil ROI. Then the neuropil DF/F was subtracted from the soma DF/F, and finally, a correction factor was added to ensure that the soma DF/F remained positive. The resulting DF/F of each cell was then deconvolved using the CaImAn package (Giovannucci et al., 2019; Friedrich et al., 2017) to obtain event rates that approximate the cell’s activity level.

#### Quantification of turn selectivity

Turn selective cells. We quantified turn-selectivity by comparing deconvolved DF/F event rates between CW and CCW trials. Turn-selectivity was then determined using the Linear-Non Linear Poisson model (see section “Statistical Model”). Cells with significance values p < 0.05 were classified as turn-selective (TS) cells.

#### Turn selectivity index

We defined this as:

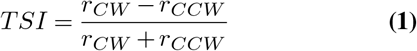

where *r* is the average event rate during CCW (or CW) trials. The turn selectivity index, *TSI*, will take on values [1, +1] and cells with *TSI* < 0 are considered CCWpreferring cells while cells with *TSI* > 0 are considered CWpreferring cells. Extreme values −1 (or +1) denote cells that fire exclusively during CCW or CW trials, respectively.

#### Spatial clustering

To determine whether turn-selective cells are spatially clustered, we plotted the difference of *TSI* for pairs of cells, ∆*TSI_ij_* = *TSI_i_ TSI_j_*, as a function of the physical distance between them. We first tested this measure using simulated FOVs containing a typical number of cells using a Gaussian mixture model (Figure S1A; MATLAB gmdistribution, two clusters with center of mass *μ* and *σ*, *μ*_1_ = [0, 0], *σ*_1_ = 1, *μ*_2_ = [1, 1], *σ*_2_ = 0.5). In the first simulated FOV, each cell is randomly assigned a *TSI* [1, +1] (to mimic a “salt-and-pepper” case). In the other simulated FOV, cells of the same cluster were assigned a similar *TSI* (to mimic the clustered case). We then averaged ∆*TSI_ij_* for each spatial distance bin (Figure S1B). For a “salt-and-pepper” organization, this measure is not dependent on the distance between cells. For a “spatially clustered” organization, this measure increases with distance. In the experimental data (example FOV in Figure S1C), we binned the spatial distance *d_ij_* from 20 μm to 400 μm into 20 bins and averaged ∆*TSI_ij_* for each bin.

## Classification of head direction tuning

### Head direction tuning curve

For each cell, these were calculated by binning

**Methods Figure 4.**
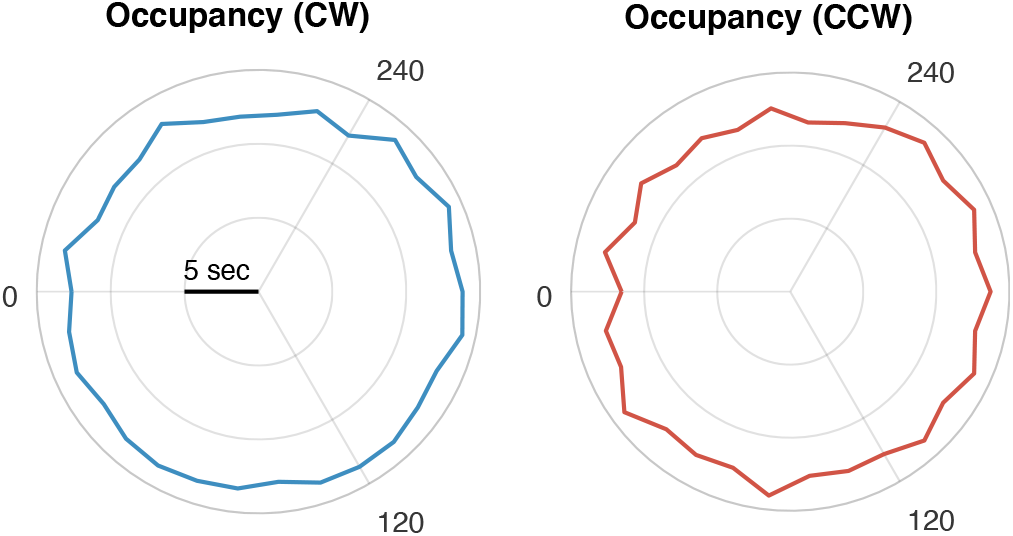
Occupancy distribution. Polar histogram of head direction occupancy during passive rotation for all CW trials (left) and all CCW trials (right)

deconvolved DF/F events in angular bins of 30°. To correct for potential sampling bias, we divided the number of events by the occupancy vector, that is, the total amount of time spent in each bin (Methods Figure 4). Lastly, the tuning curves were smoothed with a Gaussian kernel with a standard deviation of 6°.

### Classification of HD

To classify cells, we computed the mutual information denoted as *I* between HD and average event rates as in (Skaggs et al., 1993). We chose this measure because we often observed multiple peaks in the HD tuning curve. This is a more general measure than the standard mean vector length HD-score, which assumes single-peaked HD tuning curves. The mutual information of a cell is defined as:

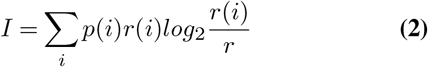

where *I* is the mutual information rate of the cell in bits per second, *i* indexes the head direction bin, *p*(*i*) is the normalized occupancy vector of the head direction, *r*(*i*) is the mean firing rate when the mouse is facing head direction *i*, and *r* is the overall mean firing rate of the cell.

We compared the mutual information to a null hypothesis determined by shuffling the data. To preserve the time dynamics of the signal, we shifted each trial by a random amount of time. We performed this operation 1,000 times to obtain a distribution of mutual information values. We defined a cell as HD-tuned if the mutual information of the original data is higher than 95 % of the mutual information in the shuffled data.

## Statistical model

We used the Linear Non-Linear Poisson (LNP) model (Hardcastle et al., 2017) to determine whether cells encode the direction and/or speed of angular head movements. We chose the LNP model because it has no prior assumptions about the tuning curve shape. In brief, these models predict the observed events by fitting a parameter to each behavioral variable. First, the probability of observing the number of events *k* for each time bin *t* is described by a Poisson process *P* (*k*(*t*) *λ*(*t*)). The estimated event rate 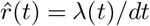, with time bin size *dt* = 32.2 ms and dimensionless quantity *λ*(*t*), is calculated as an exponential function of the weighted sum of input variables.

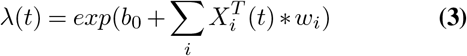

Here, *b*_0_ is a time-invariant constant, *i* indexes the behavioral variable, *X_i_* is the state matrix and *w_i_* is a vector of learned weight parameters. We designed the state matrix *X_i_* where each column is an animal state vector *x_i_* at one instant time bin *t*, defined as 1 at its current state and 0 in other states. We binned the rotation direction into two classes and the AHV into eight classes by 45°/s increments from −180°/s to +180°/s. All stationary states (0°/s) were excluded.

### Optimization of parameters

We measured performance by the log-likelihood (LLH) measure, defined as the log product of probabilities over all time bins *LLH* = log *t P* (*k*(*t*) *λ*(*t*)). The log-likelihood becomes simplified for a Poisson process:

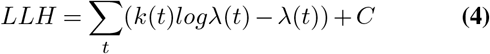

where *C* is a constant independent of the parameters *b*_0_ and *w*. To learn the parameters *w* for each cell, we maximize the log-likelihood with an additional constraint that the parameters should be smooth. That is, we find:

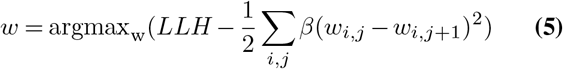

where parameter β_i_ is a smoothing hyperparameter and j cycles through adjacent components of w. Based on testing from cross-validation, we applied a uniform β = 0.01 to all cells for AHV. We optimized the parameter search using MATLAB’s fminunc function.

### Classification of tuned cells

To classify whether a cell is tuned to a specific behavioral variable, we ran two models on each cell: one without the variable, and one with the variable. If the model’s performance improved significantly with the inclusion of that variable, then the cell is classified as being tuned to the behavior. To quantify this method, we performed an 8-fold cross-validation on both models. For each fold, we used 7/8 of all trials as the training set and 1/8 of all trials as the test set. The test set LLHs of the model with the variable is compared to that of the model without the variable using paired Wilcoxon tests. Cells with a consistent increase of LLH (single tailed, p<0.05) are marked as tuned to the variable.

To estimate the percentage of classified cells expected by chance (Figure 6B), we uncoupled AHV and neural activity by randomly permuting the order of trials and following the same cell classification procedure. Averaging from a sample of 54 sessions across all brain regions and all protocols, we found that chance levels were 0.34 % for CW and CCW cells, and 0.68 % for AHV cells.

In angular velocity protocols, slow-rotating trials take longer than fast-rotating trials to complete the entire 180° rotation. This produces a bias that slow-rotating trials have a larger data sample size. To remove this bias, we truncated the data to the duration of the shortest trial.

**Methods Figure 5.**
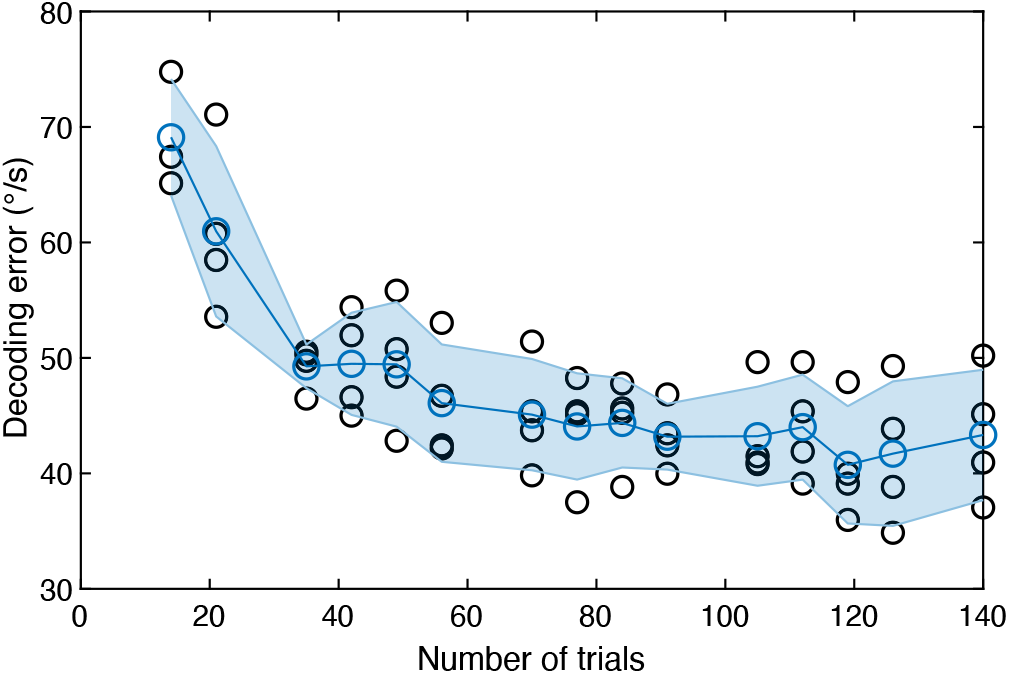
Decoding error as a function of trial number. From a single session we took an increasing number of random trials to train the model. For each number, we picked a new set of random trials and determined the decoding error. Blue line is average ± s.e.m. (4 sessions from 3 mice).

### Decoding

We used the Bayesian decoding method to reconstruct the animal’s state from its neural data. Given parameters *w*, we decode individual trials by summing the loglikelihood of observing events *k_n_*(*t*) over all cells *n* for each possible animal state. The decoded trial is the state 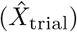 which maximizes the log-likelihood:

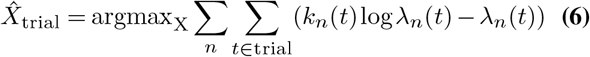

where n indexes the cell and t sums all the time bins within the trial. The decoding performance is visualized through a confusion matrix C. For the confusion matrix, each element Cij is the number of times that a trial i is decoded as trial j. The confusion matrix shows systematic errors when reconstructing the state. The confusion matrix is shown in Figure 4C. Further more, we quantified the AHV decoding error for each trial type V as the root-mean-square error:

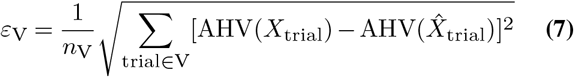

where *∊*_V_ is the decoding error, *n*_V_ is the number of trials, AHV(*X*_trial_) is the actual velocity and AHV(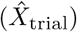) is the decoded velocity. We compared this decoding error to a null hypothesis by shuffling the trial label 1,000 times. Note that the form of the decoding error has boundary effects, i.e. the decoding error at chance level is higher for higher AHV. The overall measure of decoding error is calculated as average over all trial types.

To ensure that the experiments contain enough trials to find a reasonable decoding error, we calculated the decoding error from a random subset of trials from a control session (Methods Figure 5). Based on these data, we concluded that 40-50 rotation trials were sufficient as the decoding error did not significantly improve.

## Comparison between cortical areas and rotation conditions

We grouped the decoding performance for each brain region (V1, V2, RSC, PPC, M1+M2 and S1) and compared groups using Wilcoxon-Mann-Whitney tests. This way, we performed 15 comparative tests. Because we aimed to quantify the statistical difference between each two brain areas, we did not apply a Bonferroni correction factor to account for multiple testing. We used the same method to compare sessions under different experimental conditions (rotations in light/dark and rotations of the arena wall).

## Model of vestibular nerve activity

To simulate how mechanical activation of the vestibular organs is translated into neural impulses in the vestibular nerve (Figure S2C, F), we used a model of head motion and vestibular sensors as described in Laurens and Angelaki (2017). In brief, the model uses a Kalman filter based on a forward model of a dynamical system, defined by a set of state variables that are driven by their own dynamics, motor commands and internal or external perturbations. For a complete set of equations, we refer to Laurens and Angelaki (2017).

Two internal state variables are relevant here: the head angular velocity Ω(*t*) and hidden vestibular canal dynamics *C*(*t*). The semicircular canal signals are described by *V* (*t*) = Ω(*t*) *C*(*t*). For passive rotation, the internal state Ω(*t*) is fully described by experiment velocity profiles shown in Figure S2B, E. The canal dynamics are described as *C*(*t*) = *k*_1_*C*(*t − δt*)+*k*_2_Ω(*t*) with *k*_1_ = *τ_c_/*(1+*δt*) and *k*_2_ = *δ_t_/*(1+ *τ_c_*). We used *δt* = 32.2 ms and found the best matching fit using *τ_c_* = 2.2 s. Other constants were the default values of the model with no sensory noise. *V* (*t*) is plotted in Figure S2C and F. The code is available at: https://github.com/JeanLaurens/Laurens_Angelaki_Kalman_2017

## Fast pupil reset events

We tested whether RSC neurons respond to reafference signals generated by pupil movements, such as the fast pupil adjustments (“resets”) during a nystagmus. To determine such pupil reset events, we took the pupil position and derived the pupil velocity (Figure S3A). We then labelled pupil reset events as time points where the pupil moves at an absolute velocity over the 95th percentile. Next, we constructed the PSTH of neural activity around pupil reset events. We only used events that were not preceded by another event in a 1-second window (Figure S3C). We then averaged neural activity following a pupil reset and compared this to 1,000 instantiations of random neural activity obtained by circular time-shuffling. We classified cells as pupil reset cells if their response amplitude was larger than the 95th percentile.

To test whether the low sensitivity of the GCaMP6 indicator in response to sparse firing and the low number of pupil resets during a recording were a limiting factor in detecting pupil reset-responsive cells, we created a surrogate data set (Figure S3D). We simulated fluorescence signals of a cell that responded to pupil resets at different levels of response reliability (probability of responding to a reset) and response specificity (the overall level of background activity). First, based on previous work Sikes et al. (1988), we estimated the number of action potentials in response to a pupil reset. The resulting spike vector was then convolved with a dual exponential kernel with a rise and decay time constant of 180 and 550 ms, respectively, to match the onset and decay kinetics of GCaMP6s. Next, a vector of white noise was added to mimic realistic signal to noise levels of in-vivo fluorescence imaging. Finally, we deconvolved the DF/F of these simulated fluorescence traces, similar to what we did for experimental data.

---

**Figure S1.**
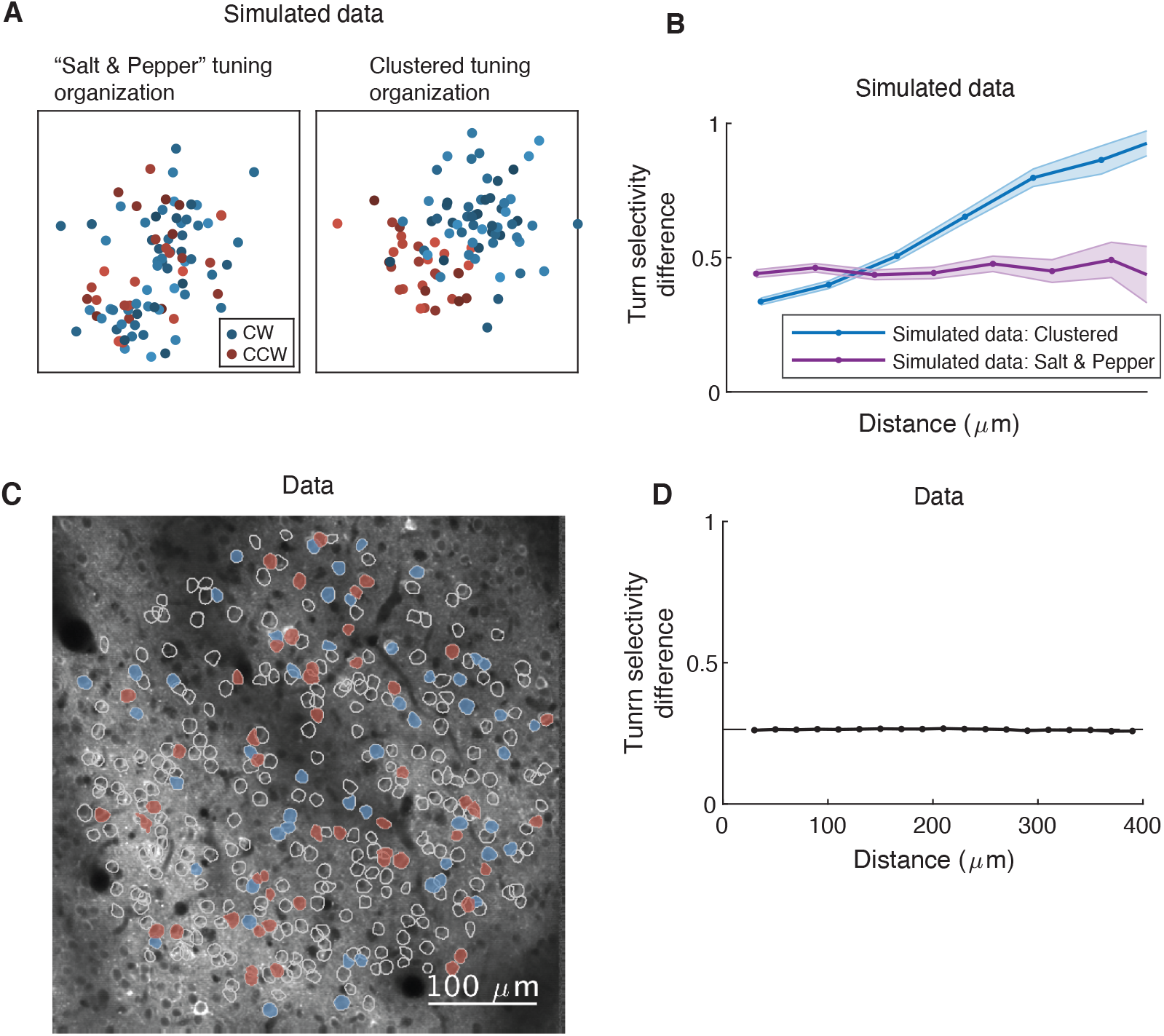
Turn-selective cells are not spatially clustered. (**A**) Simulated data showing two hypothetical forms of spatial organization between cells with similar tuning. Left: A “salt and pepper” organization lacks any spatial correlation between similarly tuned cells. Right: A “spatially clustered” organization shows neurons with similar tuning properties in closer proximity. Blue and red dots represent CW and CCW-tuned cells, respectively, on a Field of View size (FOV) that matches the typical FOV size used for two-photon imaging. (**B**) The relation between turn selectivity (CW or CCW) difference and physical distance for simulated data. The turn selectivity index difference between cells i and j defined as ∆*TSIij* = |*TSIi* − *TSIj* | is averaged for each physical distance bin in the FOV. For a "salt-and-pepper" organization, this measure is not dependent on the distance between cells. For a "spatially clustered" organization, this measure increases with distance. (**C**) Experimental data showing a two-photon microscopy image of an example FOV with turn-selective cells, color-coded in red (CCW) and blue (CW). (**D**) The relation between turn selectivity difference and physical distance for experimental data resembles more closely a "salt-and-pepper" organization (30 FOVs from 5 mice).

**Figure S2.**
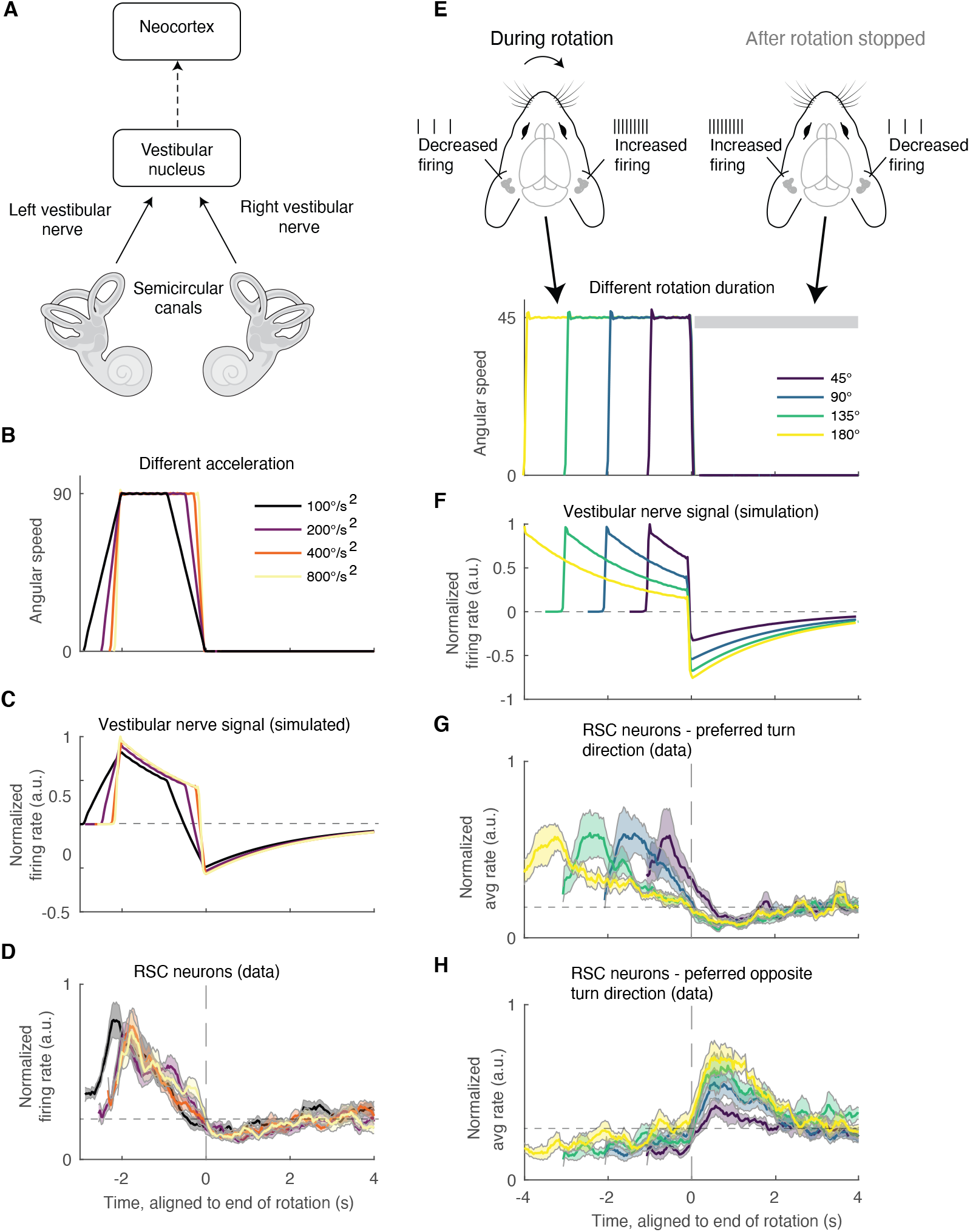
Evidence for a vestibular origin of AHV tuning in RSC. (**A**) Schematic showing a simplified pathway from the vestibular organs to the cortex. The afferent fibers of the vestibular nerve convey head motion information from the hair cells in the vestibular organs to the vestibular nucleus in the brain stem. From there on, vestibular information can take different pathways to the neocortex (dashed arrow). (**B**) Stimulation protocol: The mouse is briefly rotated at a speed of 90 deg/s using different accelerations and decelerations. (**C**) Simulations showing the response of afferent fibers in the vestibular nerve using the speed profiles in (B). (**D**) Experimental data showing the response of all turn-selective RSC neurons (average ± s.e.m DF/F event rate) using the speed profiles shown in (B) (594 turn-selective cells from 4 mice). (**E**) Stimulation protocol: The mouse is rotated at a constant speed of 45 deg/s for different durations (spanning angles of 45 to 180 degrees). Left: For brief CW rotations, fibers in the right vestibular nerve will increase firing, while those in the left vestibular nerve will decrease firing. During long rotations, however, the firing rates of both nerves will return to baseline due to the adaptation of the inertia signal in the vestibular organs. Right: When stopping the mouse after a long CW rotation, the firing rates in the left vestibular nerve will go up and the ones in the right vestibular nerve go down. This predicts that, for long CW rotations, CW-tuned cells respond during rotation, while CCW-tuned cells will respond when the rotation stops. (**F**) Simulations showing the response of afferent fibers in the vestibular nerve using the speed profiles in (E). (**G**) Experimental data showing the response of all CW-tuned neurons in RSC (average ± s.e.m DF/F event rate) using the speed profiles shown in E (330 cells from 4 mice). (**H**) Experimental data showing the response of all CCW-tuned neurons in RSC (average ± s.e.m DF/F event rate) after the rotation stops (249 cells from 4 mice).

**Figure S3.**
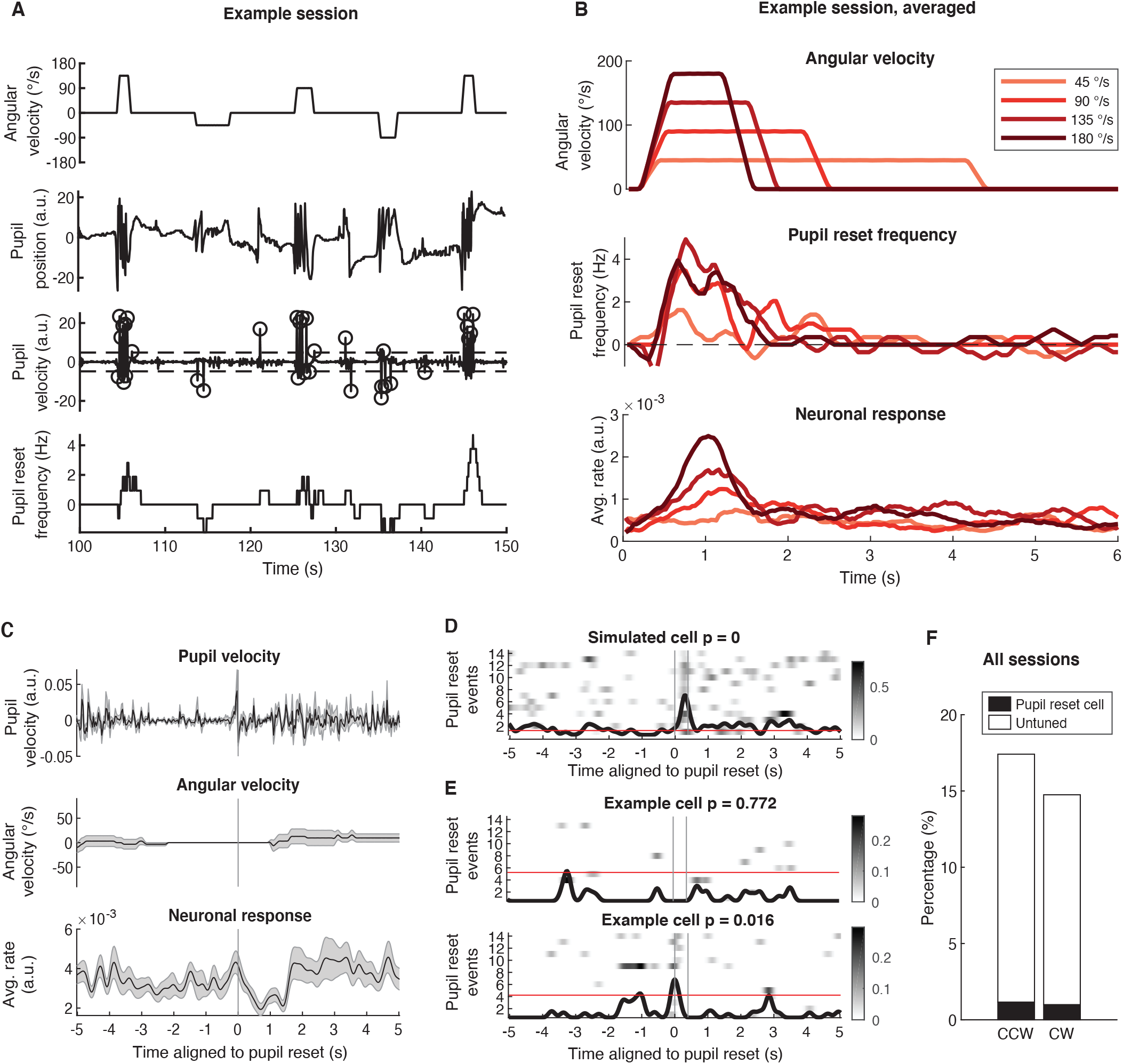
Do RSC neurons encode eye muscle reafference signals? (**A**) Example data of a mouse being rotated in CW or CCW directions at different speeds, while tracking pupil movements and neuronal activity in RSC. The experimental protocol is described in Figure 3A. 1st plot: Angular speed profile. Example sequence of 5 trials. 2nd plot: Pupil position. Note the fast pupil position resets during rotations. 3rd plot: Pupil velocity (derivative of pupil position). When the pupil velocity passed a threshold the pupil movement was defined as a fast reset event (indicated with circles). The threshold was defined as velocities exceeding the 95th percentile (two tailed distribution). 4th plot: Pupil reset events were smoothened over a 1-second window to produce the instantaneous pupil reset frequency. (**B**) Comparison of pupil reset frequency and neuronal responses in RSC, for one example session. Top: rotation velocity profile. Middle: Average pupil reset frequency. Bottom: Average response from all RSC neurons in the FOV (DF/F event rate, 287 neurons). (**C**) Because the pupil reset frequency and the angular head velocity is so correlated (shown in (B)), we also isolated fast pupil resets that occurred when the mouse was not being rotated and tried to find responsive neurons. Top, Middle: Pupil velocity and angular head velocity. Bottom: Average response of an example neuron (average ± s.e.m DF/F event rate) aligned to pupil reset. (**D**) Because of the limited number of fast pupil resets that occur outside the rotation epochs, and because of the limited sensitivity of GCaMP6s to sparse firing, we were concerned that we may not be able to detect neurons that are tuned to pupil resets. Therefore, we simulated the responses of a hypothetical pupil reset-tuned neuron using known signal to noise ratios of GCaMP6 signals during sparse firing. Panel (D) shows the average response (DF/F event rate) using only 14 pupil reset events. A neuron was deemed tuned to pupil resets when the average event rate was higher than the 95th percentile of shuffled data (red line) between 0-500 ms after the peak of the pupil reset. In case of the simulated data, the average response was well above this threshold, suggestion that, if pupil reset neurons were present in the data set, we should be able to detect them. (**E**) Experimental data showing an example neuron that did not pass the criteria, and one that may be tuned to pupil resets. (**F**) Percentage of neurons (separated into CW and CCW-tuned cells) that were significantly tuned to fast eye resets (3708 cells from 5 mice, 1.2 % cells were CW and pupil reset tuned, 0.9 % of cells were CCW and pupil reset tuned).

**Figure S4.**
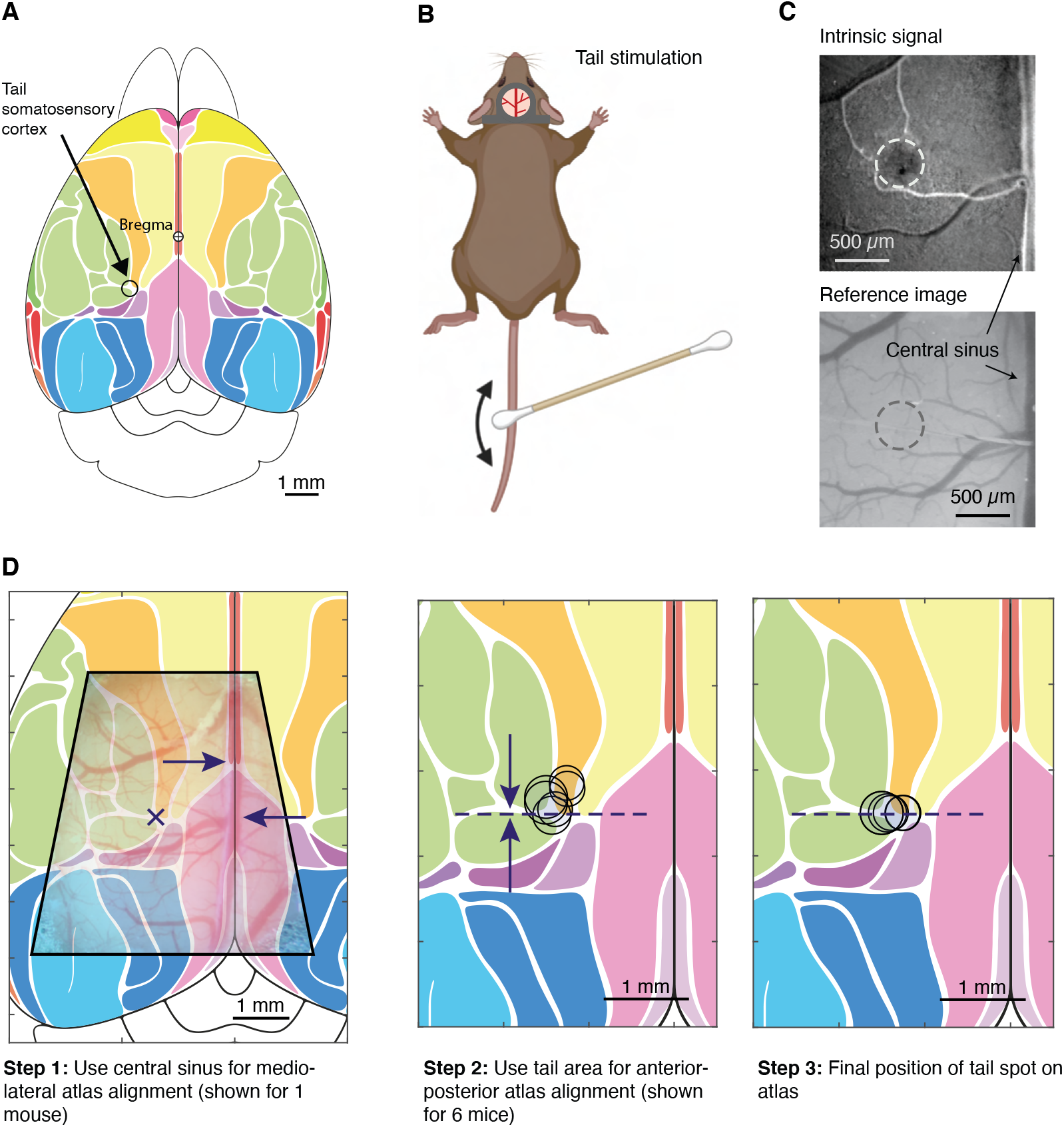
Intrinsic imaging to align two-photon maps to standardized atlas. (**A**) The tail somatosensory cortex is a small ~200-300 μm area, located at the borders of somatosensory cortex S1, primary and secondary motor cortex (S1 and S2) and posterior parietal cortex (PPC). Therefore, identifying this area in every mouse can be used to aligned two-photon microscopy-generated maps of tuned-neurons to a standardized atlas. (**B**) We used a cotton swap to stroke the tail while performing intrinsic imaging. (**C**) Top: Example of intrinsic signal response to tail stimulation. Bottom: Reference image showing blood vessel pattern. (**D**) Procedure to align maps of individual mice to standardized atlas. Left: First, the central sinus was used to align maps along the mediolateral axis. Middle: Next, the location of the tail somatosensory cortex was used to align maps along the anterior-posterior axis. Right: Final location of the tail somatosensory cortex of individual mice on the atlas.

## Supplementary Video Captions

**Video S1. Video of rotating mouse and rotating walls.** Head-restrained mouse rotated under the two-photon microscope. Example trials, rotating either the mouse or the arena wall with visual cues. Inset shows pupil tracking.

**Video S2. Video of rotating FOV and de-rotated FOV.** Example of two-photon image sequence obtained while rotating the mouse (left), and after the post-processing when images are de-rotated and corrected for brain movement (right).

**Video S3. Example of luminance artefacts during rotation under a two-photon microscope.** Even minor misalignments of the excitation laser beam will result in major luminance changes within the image during rotation.

**Video S4. Illustration of how minor excitation beam misalignment results in major luminance changes.** Left: Schematic of two-photon microscope objective and mouse brain with cranial window above dorsal (agranular) RSC. The two-photon excitation beam scans across neurons in layer 2/3. Here, for simplicity, we assume only a simple line-scan (green). We also represent inhomogeneities in the brain tissue, predominantly blood vessels, by the grey dot. Right, top: Simulated % of collected light during the line scan. Right, bottom: Simulated difference in collected light (% DF/F) when the centre ray of the excitation beam is offset from the perpendicular by 1-5°. When the centre ray of the excitation beam is perfectly perpendicular to the cranial window, there is no luminance artefact. However, if the centre ray is only 1-5° offset from the perpendicular, then luminance changes are up to 30 % DF/F.

**Video S5. Example of AHV decoding using RSC neurons.** Example trial sequence showing a mouse being rotated in the CW or CCW direction. Top: Responses of all neurons in the FOV (DF/F events). Middle: Histogram of population response (DF/F events). Bottom: Traces show the actual rotation direction and speed, and that predicted by the LNP model. For decoding, all neurons in a FOV were used regardless of whether they were turn-selective or not.

